# Thymic DC2 are heterogenous and include a novel population of transitional dendritic cells

**DOI:** 10.1101/2025.06.13.659520

**Authors:** Matouš Vobořil, Fernando Bandeira Sulczewski, Ryan J. Martinez, K. Maude Ashby, Michael Manoharan Valerio, Juliana Idoyaga, Kristin A. Hogquist

## Abstract

Myeloid cells, including dendritic cells (DCs) and macrophages, are essential for establishing central tolerance in the thymus by promoting T cell clonal deletion and regulatory T cell (Treg) generation. Previous studies suggest that the thymic DC pool consists of plasmacytoid DC (pDC), XCR1^+^ DC1 and SIRPα^+^ DC2. Yet the precise origin, development, and homeostasis, particularly of DC2, remain unresolved. Using single-cell transcriptomics and lineage-defining mouse models we identify nine major populations of thymic myeloid cells and describe their lineage identities. What was previously considered to be “DC2” is actually composed of 4 distinct cell lineages. Amongst these are monocyte-derived DCs (moDC) and monocyte derived macrophages (moMac), which are dependent on thymic interferon to upregulate MHCII and CD11c. We further demonstrate that conventional DC2 undergo intrathymic maturation through CD40 signaling. Finally, amongst “DC2” we identify a novel thymic population of CX3CR1^+^ transitional DC (tDC), which represent transendothelial DCs positioned near thymic microvessels. Together, these finding reveal the thymus as a niche for diverse, developmentally distinct myeloid cells and elucidate their specific requirements for development and maturation.

## Introduction

Thymic central tolerance is an essential process that protects the mammalian body against autoimmunity by forming a self-tolerant repertoire of T cells (Ashby and Hogquist, 2024). The scope of central tolerance is determined by the diversity of self-peptide–major histocompatibility complexes (self-p:MHC) that developing thymocytes recognize on thymic antigen presenting cells (APCs). The thymus hosts various types of APCs, including thymic epithelial cells, and a diverse spectrum of APCs of hematopoietic origin, such as dendritic cells (DCs), macrophages, and B cells (Klein and Petrozziello, 2024). The role of thymic DCs, and potentially other myeloid cells, in central tolerance was first proposed in mice, where the genetic ablation of CD11c^+^ cells led to impaired negative selection of T cells followed by the development of severe autoimmunity (Ohnmacht et al., 2009).

The thymic DC pool closely resembles the distribution of DCs in secondary lymphoid organs, including plasmacytoid DC (pDC), XCR1^+^ conventional DC type 1 (DC1), SIRPα^+^ DC2 (Li et al., 2009), as well activated DC-counterparts (aDCs), defined by CCR7 expression and increased MHC class II and co-stimulatory molecules expression (Ardouin et al., 2016). Unlike in peripheral tissues, DC activation in the thymus remains unchanged in germ-free mice, and it is not affected by genetic ablation of various innate immune signaling adaptors, indicating “sterile” forms of DC activation within the thymus (Ardouin et al., 2016; Oh et al., 2018). Recently, tonic expression of pro-inflammatory cytokines has been reported in the steady-state thymus, where they regulate the maturation and activation of various thymic DCs. For example, thymic CD301b^+^ DC2 require type II cytokines signaling (interleukin-4 (IL-4) and IL-13) for their maturation (Breed et al., 2022), while type III interferons (IFN) have been shown to regulate thymic DC1 and macrophage activation (Ashby et al., 2024). Additionally, CD40 signaling, along with the absence of single-positive (SP) thymocytes or MHC class II on hematopoietic cells results in a marked decrease in thymic aDC populations (Oh et al., 2018). Thus, tonic inflammatory signals, together with cognate interaction with SP thymocytes, promote the homeostatic activation of thymic DCs. However, the specific requirements for DC1 and DC2 maturation in the thymus remain poorly defined.

In the thymus, DC1 function is primarily associated with cross-presentation of mTEC-derived self-antigens to developing T cells facilitating the selection of regulatory T cells (Tregs) (Perry et al., 2014). Conversely, due to the high heterogeneity within SIRPα^+^ DC2, the specific function of DC2 in the thymus remains less well understood. Thymus-resident CD301b^+^ DC2 have been shown to be potent mediators of clonal deletion, as their genetic ablation alters the CD4SP thymocyte repertoire (Breed et al., 2022). Conversely, a subpopulation of SIRPα^+^ DC2 was described to originate in the periphery and thus be capable of presenting the antigens acquired in the periphery for the purpose of T cell selection (Bonasio et al., 2006). More recent research has identified the population of transendothelial DC positioned to the proximity of thymic microvessels, where they capture and present blood-born antigens to developing T cells. The positioning of these cells depends on the CX3CR1-CX3CL1 axis (Vollmann et al., 2021). Furthermore, the population of CX3CR1^+^ DCs has been observed to increase in the thymus after the weaning period of mice and these cells have been shown to be responsible for inducing the expansion of microbiota-specific T cells (Zegarra-Ruiz et al., 2021). This, along with newly described clusters of CX3CR1^+^ monocyte-derived DCs (moDC) (Vobořil et al., 2020), as well as MHC class II^+^ CX3CR1^+^ macrophages (Zhou et al., 2022), present an extraordinary challenge in elucidating the developmental and functional heterogeneity within the thymic SIRPα^+^ DC2 population.

In this study, we used single cell RNA sequencing (scRNA-seq) and various lineage-defining mouse models to investigate the origin, development and homeostasis, particularly of thymic SIRPα^+^ cells. We show that the thymic SIRPα^+^ DC population includes populations of moDC and monocyte-derived macrophages (moMac), defined by *Ms4a3^Cre^* tracing and maintained through thymic IFN signaling. We further demonstrate that DC2 undergo intrathymic activation regulated by CD40 signaling to become CCR7^+^ aDC2. Finaly, we identify a novel thymic population of transitional DC (tDC), sharing the developmental origin with pDC, that exhibit thymus-immigrating capacity and are positioned near thymic microvessels. Altogether, our study highlights the substantial heterogeneity within SIRPα^+^ DC2 compartment and provides a developmental and functional characterization of individual thymic SIRPα^+^ DC subsets.

## Results

### Single-cell RNA sequencing reveals heterogeneity in thymic DC2

To thoroughly characterize thymic DC/myeloid cell populations, we used scRNA-seq of sorted CD11c^+^ and CD11b^+^ cells from 7 week old C57BL/6 mice (**Fig. S1A**) (Breed et al., 2022). Bioinformatically, we filtered the clusters expressing Fms-related receptor tyrosine kinase 3 (*Flt3*), and colony stimulating factor 1 and 3 receptors (*Csf1r*, *Csf3r*) to identify DCs, monocyte/macrophages, and granulocytes (**Fig. S1B**). After dimensionality reduction and re-clustering, we identified 16 cell clusters based on signature gene expression (**Fig. S1C**). These clusters were assigned to 9 major populations of thymic myeloid cells by their expression of lineage-defining genes (**Fig. 1 A, B**).

**Figure 1.**
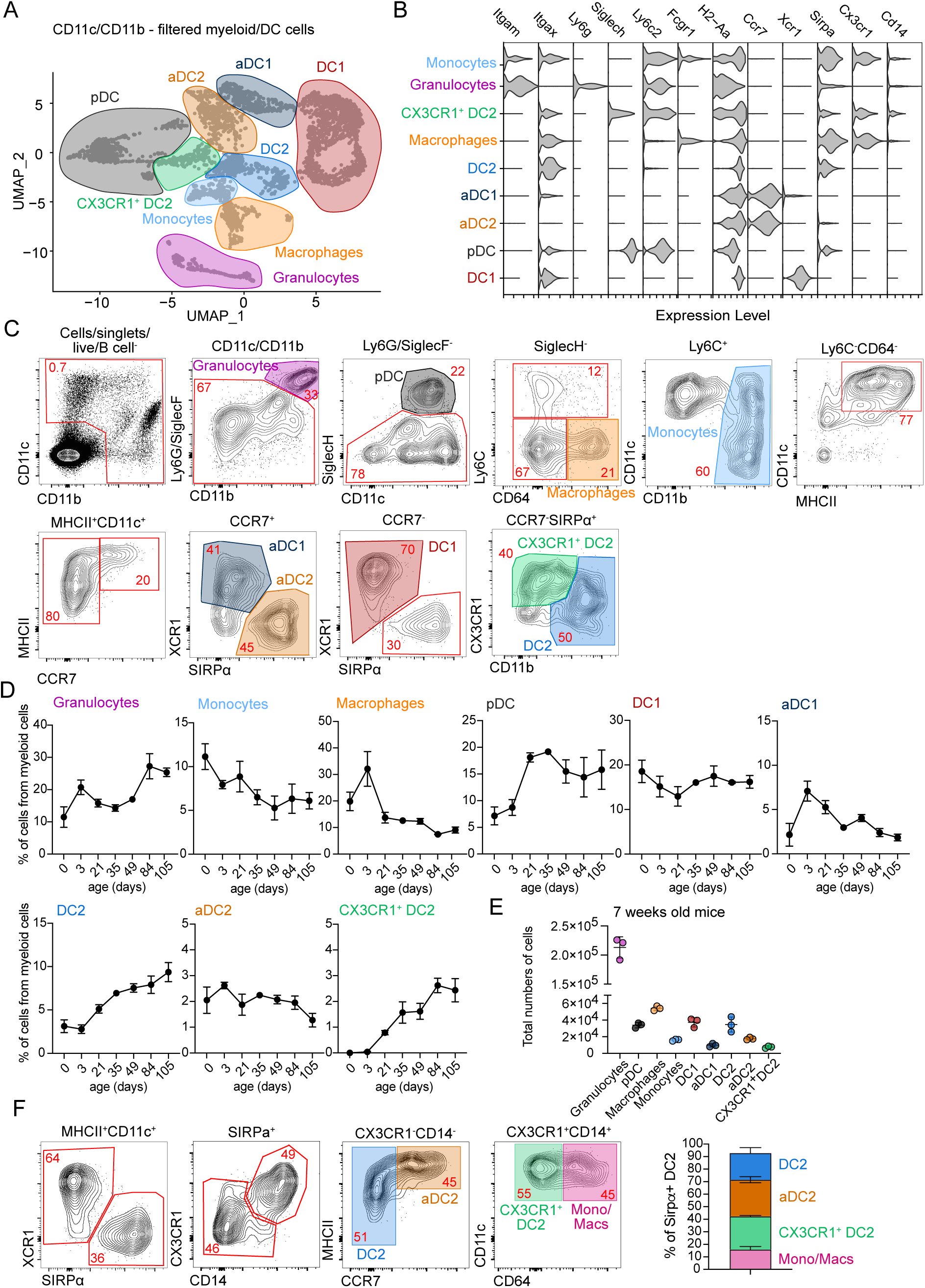
Single-cell RNA sequencing reveals heterogeneity in thymic DC2. **(A)** Single-cell RNA sequencing of CD11c^+^ and CD11b^+^ FACS-sorted cells from thymus of 7-week-old C57BL/6 mice. Cells were bioinformatically filtered to include only clusters expressing *Flt3*, *Csf1r*, and *Csf3r*. UMAP plots show the analysis of 8,514 transcriptome events, identifying 9 major clusters, marked by color-coded lines. **(B)** Violin plots displaying normalized expression of signature genes associated with cell clusters defined in (A). **(C)** Representative flow cytometry gating strategy for identifying the cell populations defined in (A) and (B). Cells were pre-gated as shown in Fig. S1D. The gating strategy identifies granulocytes (Ly6G/SiglecF^+^CD11b^+^), plasmacytoid dendritic cells (pDC; SiglecH^+^), macrophages (Ly6C^-^CD64^+^), monocytes (Ly6C^+^CD11b^+^), activated DC type I (aDC1; CD11c^+^MHCII^+^CCR7^+^XCR1^+^), aDC2 (aDC1; CD11c^+^MHCII^+^CCR7^+^SIRPα^+^), DC1 (CD11c^+^MHCII^+^ XCR1^+^), DC2 (CD11c^+^MHCII^+^SIRPα^+^CD11b^+^), and CX3CR1^+^ DC2 (CD11c^+^MHCII^+^SIRPα^+^CD11b^Low^CX3CR1^+^). **(D)** Frequency of thymic myeloid cell populations among total CD11c^+^ and CD11b^+^ cells in the thymus of C57BL/6 mice from birth (0 days old) to 105 days old mice, *n* = 2-7 mice from two independent experiments. **(E)** Total numbers of cells per thymus in 7 weeks old C57BL/6 mice, *n* = 3 mice. **(F)** Representative gating strategy for thymic CD11c^+^MHCII^+^ SIRPα^+^ cells. The graph represents the percentage distribution of individual thymic subpopulations in 7 weeks old C57BL/6 mice, *n* = 2 mice from two independent experiments. Data are shown as mean ± SD.

The thymus contains populations of granulocytes (*Csf3r, Itgam, Ly6g)*, monocytes (*Csf1r, Ly6c2, Itgam)*, and macrophages (*Csfr1, Fcgr1*). Notably, both monocytes and macrophages contain subpopulations of cells expressing higher levels of MHC class II (*H2-Aa*) and CD11c (*Itgax*), suggesting their activated phenotype (Ashby et al., 2024; Zhou et al., 2022) (**Fig 1A, B and Fig. S1C**). The thymic DC compartment includes pDC (*Siglech, Ly6c2)*, and both immature and mature conventional DC –DC1 (*Xcr1*), and DC2 (*Sirpa*) – with the mature forms defined by the expression of *Ccr7* (**Fig 1A, B**). Proliferating “cycling” subclusters, marked by *Mki67* expression, are present in both DC1 and DC2 populations (**Fig. S1C**). Interestingly, the analysis revealed a unique population of CX3CR1^+^ DC2 (in green), which clustered separately from conventional DC2 and aDC2. This population co-expressed some genes associated with pDC (*Siglech, Ly6c2)* as well as *Cd14* and *Sirpa* (**Fig 1A, B**).

We next established a flow cytometric panel enabling the discrimination of all 9 major populations of thymic myeloid cells identified by scRNA-seq (**Fig 1A, B**). To identify these cells, we gated out B cells and focused on thymic cells expressing CD11c, CD11b, or both. Granulocytes (CD11b^+^ Ly6G/SiglecF^+^), pDC (SiglecH^+^), macrophages (CD64^+^Ly6C^-^), and monocytes (Ly6C^+^ CD11b^+^) were gated sequentially (**Fig. S1D and Fig. 1C**). Within the granulocyte population we discriminated eosinophils (CD11c^+^ Ly6C^Low^) and neutrophils (CD11c^-^Ly6C^+^) (**Fig. S1E**). Thymic DC were defined as CD64^-^Ly6C^-^ CD11c^+^MHCII^+^, and activated DCs were discriminated as MHCII^High^ CCR7^+^. The expression of XCR1 and SIRPα was used to identify cells of DC1 and DC2 lineage, respectively. Notably, CX3CR1^+^ DC2 did not express the conventional DC1 marker XCR1 or the DC2/monocyte marker CD11b (**Fig. 1C**).

To evaluate age-related changes in the thymic myeloid cell compartment, we applied the previously described gating strategy and defined the proportion of individual subsets within the CD11c/CD11b population from mice aged 0 to 105 days (15 weeks). Consistent with prior findings, the DC1 and granulocyte populations maintained stable proportions from birth, whereas DC2 and pDC numbers increased during the first weeks of life (**Fig. 1D**) (Breed et al., 2022). The aDC1, aDC2, and macrophage populations peaked around 3 days of life and subsequently decline with age. Interestingly, the CX3CR1^+^DC2 population was absent in the thymus before 21 days of age, suggesting these cells may correspond to previously described thymus-immigrating CX3CR1^+^ DCs (Zegarra-Ruiz et al., 2021) (**Fig. 1D**). Numerically, by 6 weeks of age, eosinophils were the most abundant thymic myeloid cells followed by macrophages (**Fig. S1E and Fig. 1E)**. pDC, DC1, and DC2 constitute nearly equal fractions of thymic cells, while aDC1, aDC2, monocytes, and CX3CR1^+^ DC2 remained rare (**Fig. 1E)**.

Based on the above data, the population of thymic MHCII^+^CD11c^+^SIRPα^+^ cells, originally identified as conventional DC2, shows much higher internal heterogeneity (**Fig. 1B, C**). Previously, it was described that SIRPα^+^ DC contain a population of monocyte derived DC (moDC) defined by the expression of CX3CR1, CD14, and Ly6C (Vobořil et al., 2020). Here, transcriptional data suggest that, in addition to moDC, thymic SIRPα^+^ DC also contain a distinct subpopulation of CX3CR1^+^ DC2 that cluster separately from monocytes and macrophages (**Fig 1A, B**). Flow cytometry analysis confirmed this, showing that thymic SIRPα^+^ DC contain 4 subsets: conventional DC2, aDC2, CX3CR1^+^ DC2, and CD64^+^ monocyte/macrophage-derived cells, with all populations present in roughly similar proportion (**Fig. 1F**).

### The thymus contains interferon-activated populations of monocyte-derived DC and macrophages

As described above, the thymus contains a primary population of monocytes (Ly6C^+^CD11b^+^) and macrophages (CD64^+^), of which some upregulate CD11c, MHCII, and SIRPα, which are classic markers of conventional DC2 (**Fig. 1B, C**). Thus, cells with monocyte and macrophage markers constitute approximately 15% of the thymic conventional DC2 gate (**Fig. 1F**). Therefore, accurately distinguishing thymic monocyte/macrophage populations is essential for understanding the true heterogeneity of DC2s. Thymic macrophages are known to consist of two major subpopulations, defined by the expression of *Timd4* and *Cx3cr1* (Zhou et al., 2022), and depend on transcriptional factor *Nr4a1* (Tacke et al., 2015). It has been reported that *Timd4^+^* macrophages are of embryonic origin, while *Cx3cr1^+^* macrophages are derived from adult hematopoietic stem cells (Zhou et al., 2022).

To comprehensively analyze the heterogeneity in thymic monocyte/macrophage populations, we bioinformatically filtered cells expressing the transcription factor *Mafb*, which distinguishes monocytes and macrophages from other immune lineages (Wu et al., 2016) (**Fig. 2A)**. Re-clustering the data using only *Mafb*^+^ cells revealed 5 major subpopulations of thymic monocytes and macrophages. These included classical monocytes (*Ly6c2*, *Itgam*, *Cx3cr1*), classical macrophages (*Fcgr1*, *Adgre1*, *Mertk*, *Timd4*) and a population we will refer to as “monocyte derived DC” (moDC) (*Ly6c2*, *Fcgr1, Cx3cr1*, *H2-Aa*, *Itgax*), and “monocyte-derived macrophages” (moMac), defined by the expression of macrophage markers (*Adgre1*, *Mertk, Vcam1).* Consistent with previous findings (Zhou et al., 2022), thymic moMac exhibited a subset of proliferating cells, which we refer to as the *Mki67*^+^ “cycling” macrophage subpopulation. Finally, the thymus also harbors a small population of LYVE-1^+^ macrophages, characterized by the expression of *Lyve1* and *C5ar1* (CD88) (**Fig. 2B, C**).

**Figure 2.**
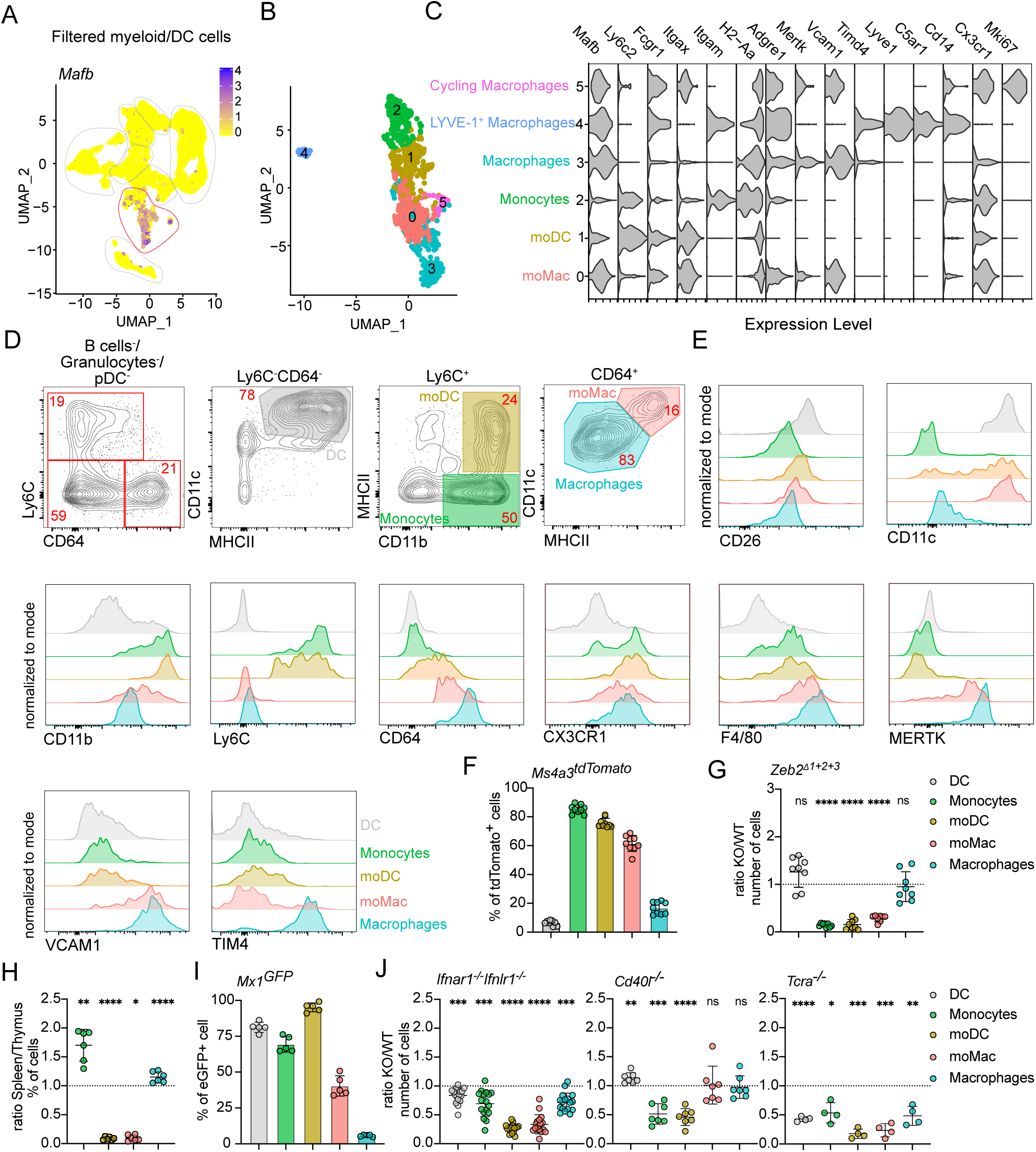
The thymus contains interferon-activated populations of monocyte-derived DC and macrophages. **(A)** Feature plot showing the normalized expression of *Mafb* in clusters identified in Fig. 1A. **(B)** Single-cell RNA sequencing of CD11c^+^ and CD11b^+^ FACS-sorted cells from thymus of 7-week-old C57BL/6 mice. Cells were bioinformatically filtered to include only clusters expressing *Mafb*. UMAP plots show the analysis of 1,020 transcriptome events, identifying 5 clusters. **(C)** Violin plots displaying the normalized expression of signature genes associated with cell clusters defined in (B). **(D)** Representative flow cytometry gating strategy for identifying the four major populations defined in (B) and (C). The gating strategy identifies monocytes (Ly6C^+^CD11b^+^MHCII^-^), monocyte-derived DC (moDC; Ly6C^+^CD11b^+^MHCII^+^), macrophages (CD64^+^CD11c^-^MHCII^-^), and monocyte-derived macrophages (moMac; CD64^+^CD11c^+^MHCII^+^). **(E)** Representative flow cytometry plots showing normalized expression of CD26, CD11c, CD11b, LY6C, CD64, CX3CR1, F4/80, MERTK, VCAM1, and TIM4 in thymic dendritic cells (DC; Ly6C^-^CD64^-^CD11c^+^MHCII^+^) and thymic monocyte and macrophage populations described in (D). **(F)** Frequency of tdTomato^+^ thymic cells (gated as in D) in *Ms4a3^Cre^* ROSA26^tdTomato^ (*Ms4a3^tdTomato^*) mice, *n* = 9 mice from 3 independent experiments. **(G)** Numbers of thymic DC, monocyte, and macrophage (gated as in D) populations in *Zeb2^Δ1+2+3^* mice, shown as KO/WT ratio of cell numbers, *n* = 8 mice from 3 independent experiments. **(H)** Numbers of thymic and splenic cells (gated as in D), shown as spleen/thymus ratio of cell frequencies, *n* = 6 mice from 3 independent experiments. **(I)** Frequency of eGFP^+^ thymic cells (gated as in D) in *Mx1^eGFP^* mice, *n* = 5 mice from 2 independent experiments. **(J)** Numbers of thymic cells (gated as in D) in *Ifnar1^-/-^Ifnlr1^-/-^, Cd40l^-/-^*, and *Tcra^-/-^* mice, shown as KO/WT ratio of cell numbers, *n* = 4-17 mice from at least 2 independent experiments. Data are shown as mean ± SD. Statistical analysis was performed by a one-sample *t* test and Wilcoxon test with a theoretical mean of 1, \**p*≤0.05, \*\**p*≤0.01, \*\*\**p*≤0.001, \*\*\*\**p*≤0.0001, n.s. = not significant.

Using transcriptional data (**Fig. S2A**), we designed a flow cytometry panel to define the major populations: monocytes (Ly6C^+^CD11b^+^MHCII^-^), macrophages (Ly6C^-^CD64^+^MHCII^Low^CD11c^Low^), moDC (Ly6C^+^CD11b^+^MHCII^+^), and moMac (Ly6C^-^CD64^+^MHCII^High^CD11c^High^). Notably, moDC also expressed CD64, although their expression levels were lower than in moMac or classical macrophages (**Fig. 1F and Fig. 2E**). We also verified the presence of LYVE-1^+^ macrophages in the thymus, using CD88 staining and testing additional markers. These cells are very rare as they constitute approximately 1.5% of all thymic macrophages (**Fig. S2B**). Consistent with their expression of myeloid lineage restricted *Mafb*, these populations expressed only very low levels of DC-defining molecules CD26 or *Flt3*, suggesting minimal DC contamination in the gating strategy (**Fig. 2E, S1B**). To validate our gating strategy, we tested the expression of several monocyte/macrophage prototypical markers via flow cytometry. Our results indicate that the macrophage population corresponds to the previously described *Timd4*^+^ macrophages (Zhou et al., 2022), as these cells upregulate MERTK and TIM4 while exhibiting lower CD11b expression (**Fig. 2E**). Despite the transcriptional similarities between moDCs and moMacs, the upregulation of macrophage-specific markers (MERTK, VCAM1) in moMacs further supports their classification as part of the macrophage lineage (**Fig. 2E**).

Although conventional DC2 and moDC/moMac exhibit distinct transcriptional profiles, establishing the lineage origin of these cells based on surface markers alone remains challenging. To verify the monocyte-origin of moDC and moMac we utilized the *Ms4a3^Cre^* ROSA26^tdTomato^ (*Ms4a3^tdTomato^*) fate-mapping system. The *Ms4a3* gene is specifically expressed in granulocyte-monocytes progenitors (GMP), but not in monocyte-DC progenitors (MDP) or subsequent DC progeny, enabling clear tracking of monocyte-derived cells (Liu et al., 2019). Our data indicate high levels of recombination within the monocyte, moDC, and moMac populations, strongly supporting their monocyte-origin (**Fig. 2F**). Conversely, the conventional DC population showed minimal recombination, confirming the very limited contamination by DCs in our gating strategy. Interestingly, the macrophage population exhibited low recombination levels as well, suggesting that these cells predominantly represent embryonically derived tissue-resident macrophages (**Fig. 2F**) (Liu et al., 2019). These findings were corroborated by using mice carrying triple mutations in *Zeb2* enhancer (*Zeb2^Δ1+2+3^*), which results in the absence of conventional DC2 and monocytes but not tissue-resident macrophages (Liu et al., 2022). In this system we observed the depletion of monocytes, moDC, and moMacs, while macrophages remained unaffected (**Fig. 2G**).

Both moDCs and moMacs resemble fully matured APC populations, characterized by high expression of MHCII and various co-stimulatory molecules (**Fig. S2C**). To explore their distribution in peripheral tissues, we compared these populations in the thymus and spleen. Surprisingly, moDCs and moMacs were almost entirely absent in the steady-state spleen, unlike in the thymus (**Fig. 2H, S2D**), suggesting that their maturation occurs intrathymically. Previously, it was reported that most thymic APCs respond to sterile IFN at steady state, evidenced by upregulating *Mx1* (Ashby et al., 2024). We further tested monocytes and macrophages for their ability to respond to thymic IFN and examined their dependency on IFN signaling. The analysis of *Mx1^eGFP^*mice confirmed that monocytes, moDC and 50% of moMAC express GFP, whereas classical macrophages do not (**Fig. 2I**). We hypothesize that macrophages are localized primarily in the thymic cortex (Zhou et al., 2022), whereas IFN production was found in the medulla (Ashby et al., 2024). Consistent with this, moDC and moMac maturation depended on thymic IFNs as mice lacking type I and III IFN receptors (*Ifnar1^-/-^Ifnlr1^-/-^)* exhibited a significant reduction in these populations (**Fig. 2J**). Notably, while moMac maturation depended largely on type III IFNs, moDC maturation depended on both types of IFNs (**Fig. S2E).** Given the role of CD40 signaling and SP thymocytes in thymic DC maturation (Oh et al., 2018), we tested their involvement in monocyte and macrophage maturation. Interestingly, CD40 signaling appeared to play a minimal role, as *Cd40l^-/-^* mice showed only a minor reduction in total monocyte and moDC numbers, with no specific impact on moMacs. Conversely, the absence of SP thymocytes in *Tcra^-/-^* mice significantly reduced DC, moDC and moMac populations, while other cell types, such as granulocytes and pDC, remained unaffected (**Fig. 2J**). This finding highlights the key role of SP thymocytes in monocyte maturation, through driving IFN production in TEC.

Together, the thymus contains populations of monocytes and macrophages, along with monocyte-derived moDCs and moMacs, whose maturation relies on IFN signaling and the presence of SP thymocytes.

### Activation of conventional DC1 and DC2 requires distinct signals

After characterizing thymic populations of moDCs and moMacs, we focused on the heterogenous populations of thymic DCs. To distinguish DCs from other lineages, we bioinformatically filtered cells expressing *Flt3*. Re-clustering the data reveled 11 subclusters of *Flt3*^+^ cells (**Fig. S3A, B**), which we further classified into 7 major thymic DC clusters (**Fig. 3A, B**). Within these, we annotated the populations of pDCs, DC1, and DC2, all of which included cycling *Mki67*-expression cells, as well as a distinct subset of CX3CR1^+^ DC2 (**Fig. 3A an Fig. S3A**).

**Figure 3.**
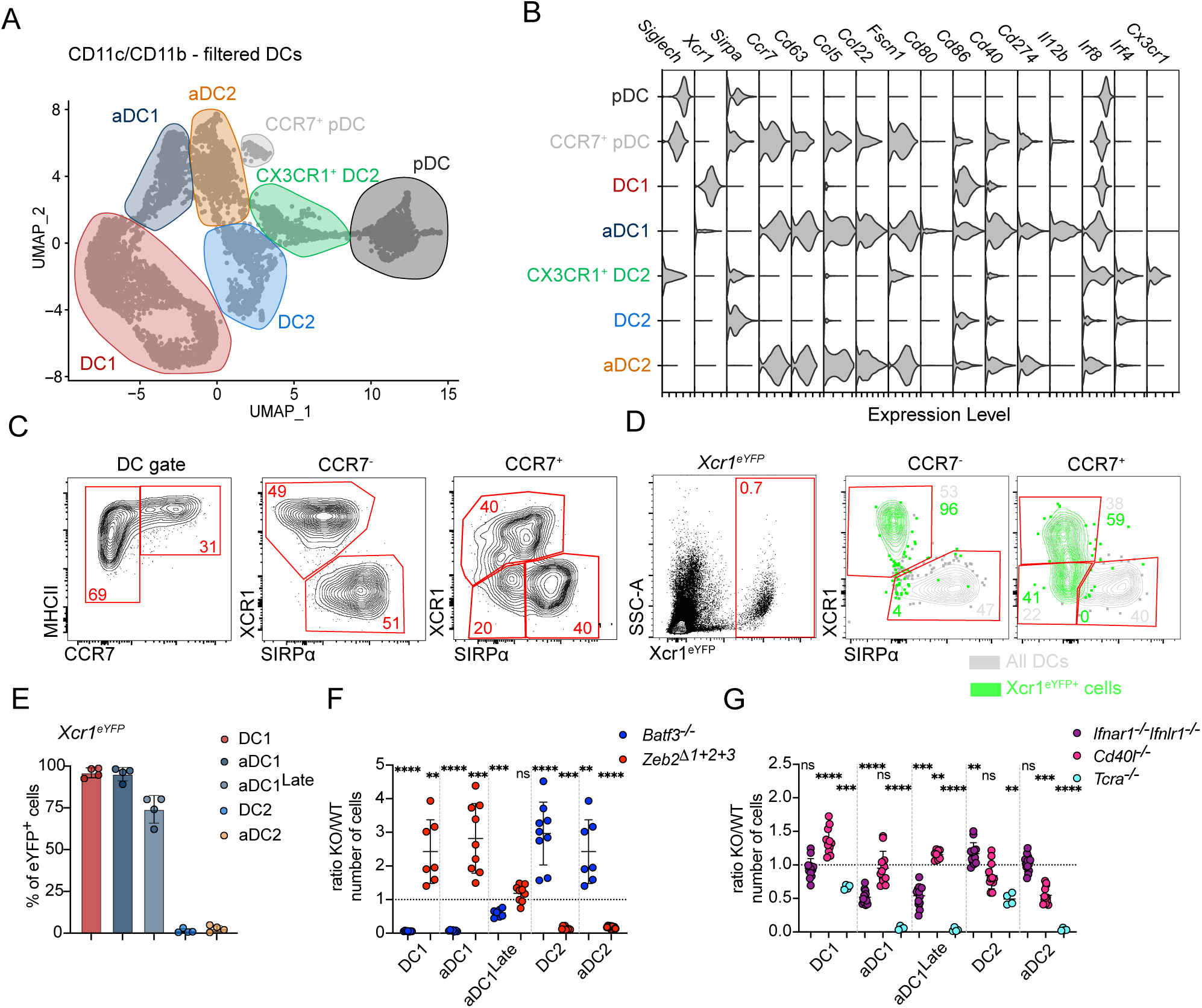
Activation of conventional DC1 and DC2 requires distinct signals. **(A)** Single-cell RNA sequencing of CD11c^+^ and CD11b^+^ FACS-sorted cells from thymus of 7-week-old C57BL/6 mice. Cells were bioinformatically filtered to include only clusters expressing *Flt3*. UMAP plots show the analysis of 6,928 transcriptome events, identifying 7 major clusters, marked by color-coded lines. **(B)** Violin plots displaying the normalized expression of signature genes associated with cell clusters defined in (A). **(C)** Representative flow cytometry gating strategy for identifying thymic dendritic cells (DC; Ly6C^-^CD64^-^ CD11c^+^MHCII^+^), DC1 (CCR7^-^XCR1^+^), activated DC1 (aDC1; CCR7^+^XCR1^+^), DC2 (CCR7^-^SIRPα^+^), and aDC2 (CCR7^+^SIRPα^+^). **(D)** Representative flow cytometry plots showing expression of eYFP by thymic DC populations described in (C) in *Xcr1^iCre^Rosa26^eYFP^* (*Xcr1^eYFP^*) mice. Gray cells represent all thymic cells gated as in (C); green cells represent eYFP^+^ cells. **(E)** Frequency of eYFP^+^ cells (gated as in C) in *Xcr1^iCre^Rosa26^eYFP^* (*Xcr1^eYFP^*) mice, *n* = 4 mice from 3 independent experiments. **(F)** Numbers of thymic DCs (gated as in C) in *Batf3^-/-^* and *Zeb2^Δ1+2+3^* mice, shown as KO/WT ratio of cell numbers, *n* = 7-9 mice from 3 independent experiments. **(G)** Numbers of thymic DCs (gated as in C) in *Ifnar1^-/-^Ifnlr1^-/-^, Cd40l^-/-^*, and *Tcra^-/-^* mice, shown as KO/WT ratio of cell numbers, *n* = 4-13 mice from at least 2 independent experiments. Data are shown as mean ± SD. Statistical analysis was performed by a one-sample *t* test and Wilcoxon test with a theoretical mean of 1, \*\**p*≤0.01, \*\*\**p*≤0.001, \*\*\*\**p*≤0.0001, n.s. = not significant.

We identified three populations of *Ccr7*^+^ cells corresponding to the previously defined mature population of activated DCs (aDCs). According to an earlier study, activated DC1 have undergone homeostatic maturation within the thymus, characterized by the overexpression of “MAT ON” genes, including chemokines, cytokines, and costimulatory molecules (Ccr7, *Ccl5*, *Ccl22*, *Fscn1*, *Cd40*, and *Il12b*) (Ardouin et al., 2016). The expression of these genes was significantly enriched in the cluster annotated as aDC1 confirming their fully activated phenotype (**Fig. 3B**). Interestingly, we also detected the expression of “MAT ON” gene expression in a population of CCR7^+^ DC2 and CCR7^+^ pDC, suggesting they may represent thymus-specific subsets of homeostatically activated DC2 and pDC, respectively (**Fig. 3A, B**). Flow cytometry analysis further confirmed the expression of CCR7 on thymic DC2 and pDC, revealing that approximately 45% of DC2 and 7% of thymic pDC represent the activated state (**Fig. 3D and Fig. S3C**). Conversely, XCR1 staining did not validate the presence of *Xcr1^+^* pDC identified through scRNA-seq analysis so these cells were excluded from analysis (**Fig. S1C and S3C**). After gating out pDC, antibody staining of MHCII^High^ CCR7^+^ DCs distinguishes the total aDC population, with XCR1 and SIPRα protein staining identifying aDC1 and aDC2, respectively. The expression of co-stimulatory molecules CD80, CD86, CD40, PD-L1 and CD63 clearly confirms the activated phenotype of both populations (**Fig. S3D**). Notably, the protein expression of XCR1 and SIPRα in CCR7^+^ populations is lower than in their immature CCR7^-^ counterparts (**Fig. 3C**). This corresponds to the almost negligible mRNA expression of *Xcr1* and *Sirpa* in these aDCs, making them much harder to discriminate by scRNA-seq (**Fig. S3E**). Furthermore, the flow cytometry analysis of MHCII^High^ CCR7^+^ reveals a subpopulation of XCR1^-^SIPRα^-^ double-negative cells, whose lineage-specific origin we wanted to determine (**Fig. 3C**).

To determine the origin of double-negative aDCs, we used *Xcr1^iCre^Rosa26^eYFP^* DC1 lineage-tracing mouse model, tracking the history of *Xcr1* expression (Ferris et al., 2020). As expected, eYFP expression was restricted to DC1 and aDC1 cells but was also highly enriched in XCR1^-^SIPRα^-^ double-negative aDCs, with approximately 75% of these cells express eYFP despite the complete absence of XCR1 protein expression (**Fig. 3D, E**). Moreover, eYFP^+^ cells with low or negligible XCR1 expression showed an increased proportion of CCR7^+^ cells, suggesting that DC1 downregulate XCR1 upon activation while acquiring the CCR7^+^ aDC phenotype (**Fig. S3F**). Based on this, XCR1^-^SIPRα^-^ double-negative aDCs likely represent a later stage of DC1 activation (aDC1^Late^), corresponding to the previously identified human aDC3 cells (Park et al., 2020).

Whereas the origin of aDC1 has been attributed to DC1-lineage (Ardouin et al., 2016), the ontogeny of aDC2 has not yet been reported. To investigate this further, we used two lineage-depleting mouse models: *Batf3^-/-^* mice, which lack DC1 cells (Hildner et al., 2008), and mice carrying triple mutations in the *Zeb2* enhancer (*Zeb2^Δ1+2+3^*), which lack DC2 lineage (Liu et al., 2022). Analysis of thymic CCR7^+^ aDCs from these mice showed that both XCR1^+^ aDC1 and XCR1^-^ aDC1^Late^ cells were significantly reduced only in *Batf3^-/-^* mice, whereas SIRPα^+^ aDC2 were depleted in *Zeb2^Δ1+2+3^* mice (**Fig. 3F and S3G**). This clearly confirms that despite their extensive transcriptional similarities, aDC1 and aDC2 originate from distinct precursors. Interestingly, the total number of thymic DCs remains unchanged in both *Batf3^-/-^* and *Zeb2^Δ1+2+3^*, as the depletion of one DC subset is compensated by an increase in the other subset (**Fig. 3F**).

The specific requirements for DC1 and DC2 maturation in the thymus remain poorly defined. Previous studies suggested that the maturation of both thymic DC1 and DC2 is markedly reduced in mice lacking CD40 signaling, as well as in those lacking single-positive thymocytes (Oh et al., 2018). More recently, type I and type III IFNs were shown to regulate maturation of DC1 but not DC2, despite both populations expressing IFN receptors (Ashby et al., 2024). We applied the previously described gating strategy to verify the dependence of aDC1 populations, as well as aDC2 on IFNs. As expected, the numbers of aDC1 and aDC1^Late^ cells were significantly reduced in *Ifnar1^-/-^Ifnlr1^-/-^*mice, whereases aDC2 remained unchanged (**Fig. 3G**). As previously described, this reduction was primarily associated with diminished type III IFN signaling (**Fig. S3H**). Conversely, mice deficient for CD40-ligand (*Cd40l^-/-^*) showed a significant reduction in aDC2 numbers, but no decrease in either aDC1 or aDC1^Late^ cells (**Fig. 3G**). These findings indicate that despite their transcriptional similarities, aDC1 and aDC2 require distinct signals for thymic maturation. Moreover, *Tcra^-/-^* mice showed significantly altered activation of both DC1 and DC2 populations, highlighting the role of single-positive thymocytes in DC maturation–either by providing CD40L or driving IFN production (**Fig. 3G**).

Together, these finding demonstrate that the thymus harbors fully activated subsets of DC1 and DC2, whose maturation depends on IFN signaling and CD40L signaling, respectively.

### The thymus contains a population of CX3CR1^+^ transitional DC (tDC)

Apart from moDC and moMacs, thymic CX3CR1^+^SIRPα^+^ DCs also include a distinct subpopulation of CX3CR1^+^ DC2, which clusters separately from conventional DC2 and aDC2 (in green, **Fig. 1A, F**). Based on their transcriptional profile, CX3CR1^+^ cells share similarity to DC2 but also to thymic pDCs (**Fig. 3A, B**). To investigate this further, we compared the transcriptional profiles of thymic DC2, CX3CR1^+^ DC2, and pDCs by bioinformatically filtering and re-clustering these populations (**Fig. 4A, B**). This analysis revealed that CX3CR1^+^ DC2 express some genes characteristic of pDCs–such as *Ly6c2*, *Siglech*, *Ly6d* and *Tcf4*–as well as some characteristic of DC2, including *Irf4*, *Cd209a*, *Klf4* and *Mgl2*. Additionally, we identified a set of genes uniquely expressed in the CX3CR1^+^ DC2 population, such as *Cd209e*, *Ngfr, Cx3cr1* and *Cd14* (**Fig. 4C**). Flow cytometric analysis confirmed the expression of TCF4, CX3CR1, and CD14 at the protein level, defining this population as TCF4^+^CX3CR1^+^CD14^+^ and SiglecH^-^CD11b^Low^ (**Fig. 4D**). This pattern suggested to us that thymic CX3CR1^+^ DC2 might represent a population called transitional DCs (tDC), recently described in peripheral tissues (Leylek et al., 2019; Sulczewski et al., 2023; Rodrigues et al., 2023). tDC were initially identified in human blood and have been shown to be conserved between mice and humans (Alcántara-Hernández et al., 2017; Leylek et al., 2019; Villani et al., 2017). The term “transitional DC” was used to highlight their transcriptomics, phenotypic and functional features, which span characteristics of both pDCs and DC2 (Leylek et al., 2019). Therefore, we decided to employ lineage marking and fate mapping approaches to test if thymic “CX3CR1^+^ DC2” are equivalent to peripheral tDC.

**Figure 4.**
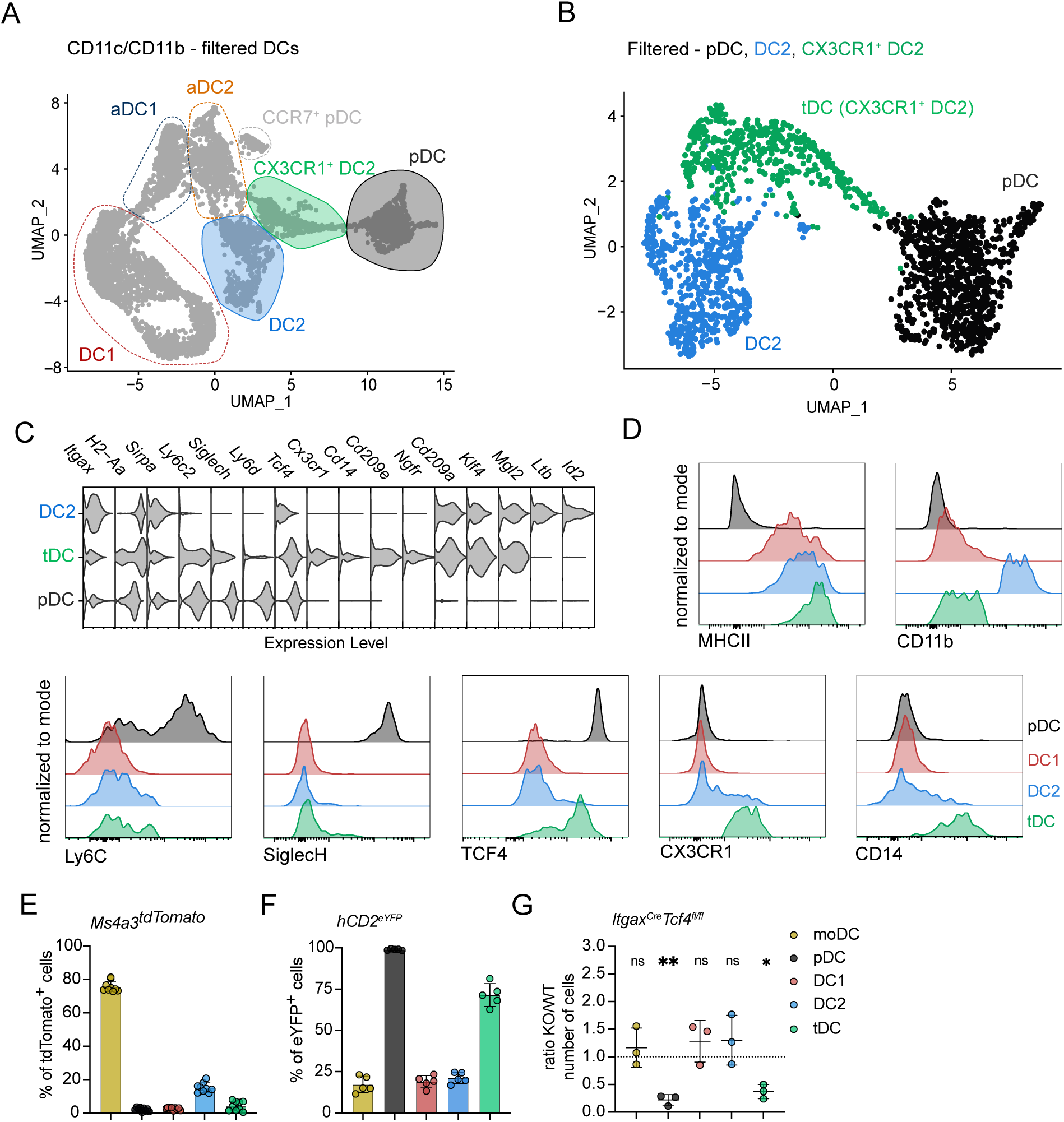
The thymus contains a population of CX3CR1^+^ transitional DC. **(A)** Single-cell RNA sequencing of CD11c^+^ and CD11b^+^ FACS-sorted cells from thymus of 7-week-old C57BL/6 mice. Cells were bioinformatically filtered and displayed as described in Fig. 3A. DC2, CX3CR1^+^ DC2, and pDC are marked by color-coded lines. **(B)** UMAP plots showing the distribution of filtered DC2, CX3CR1^+^ DC2, and pDC thymic populations defined in A. **(C)** Violin plots displaying the normalized expression of signature genes associated with cell clusters defined in (B). **(D)** Representative flow cytometry plots showing the normalized expression of MHCII, CD11b, Ly6C, SiglecH, TCF4, CX3CR1, and CD14, in thymic dendritic cells populations defined in (B) and gated according to the Fig. S4B. **(E)** Frequency of tdTomato^+^ thymic dendritic cells populations (gated as in Fig. S4B) in *Ms4a3^Cre^*ROSA26^tdTomato^ (*Ms4a3^tdTomato^*) mice, *n* = 9 mice from 3 independent experiments and moDC are used as control. **(F)** Frequency of eYFP^+^ thymic DC populations (gated as in Fig. S4B) in *hCD2^iCre^*ROSA26^eYFP^ (*hCD2^eYFP^*) mice, *n* = 5 mice from 3 independent experiments. **(G)** Numbers of thymic populations (gated as in Fig. S4B) in *Itgax^Cre^Tcf4^fl/fl^* mice, shown as KO/WT ratio of cell numbers, *n* = 3 mice from 2 independent experiments, *Itgax^Cre-^* mice were used as controls. Data are shown as mean ± SD. Statistical analysis was performed by a one-sample *t* test and Wilcoxon test with a theoretical mean of 1, \**p*≤0.05, \*\**p*≤0.01, n.s. = not significant.

Phenotypically, CX3CR1^+^ DC2 resemble monocyte-derived cells by the expression of CX3CR1 and CD14; however, tDC are not of monocyte origin but instead share a developmental lineage with pDCs (Sulczewski et al., 2023; Rodrigues et al., 2024). To investigate the origin of CX3CR1^+^ DC2, we assessed their recombination levels in the monocyte lineage-tracer *Ms4a3^Cre^* ROSA26^tdTomato^ mice. Indeed, CX3CR1^+^ DC2 were not marked by tdTomato in these mice (**Fig. 4E**), indicating that they are not a monocyte-derived population. Alternatively, we utilized a pDC-specific lineage-tracing model, expressing iCre under the human CD2 promoter (*hCD2^iCre^*), crossed with *ROSA26^eYFP^* mice (*hCD2^eYFP^*) (Siegemund et al., 2015; Dress et al., 2019; Sulczewski et al., 2023). In these mice, eYFP labeling was detected in both thymic pDCs and tDC populations, whereas other thymic DC populations showed low levels of YFP (**Fig. 4F**). Finally, we used *Itgax^Cre^Tcf4^fl/fl^* mice, which lack TCF4 expression specifically in CD11c-expressing cells. TCF4 has been previously described as essential for pDC and tDC development (Cisse et al., 2008; Sulczewski et al., 2023). Genetic ablation of TCF4 significantly reduced the numbers of both thymic pDC and tDC populations (**Fig. 4G**), confirming their shared developmental origin.

Overall, the distinct transcriptional profile of CX3CR1^+^ DC2, their shared origin with pDCs, and their developmental dependence on TCF4 expression clearly identify these cells as previously unrecognized thymic tDCs.

### tDC represent transendothelial cells

Thymic tDCs represent a mature population of DCs, expressing high expression of MHCII, comparable to that of CCR7^+^ aDCs (**Fig. S4A**). Additionally, tDCs express various co-stimulatory molecules, albeit at lower levels than conventional aDCs (**Fig. S4A**). This suggests that, while thymic tDC are activated cells, their activation state remains distinct from conventional thymic DCs. To investigate their mode of activation, we quantified the numbers of thymic tDCs in *Ifnar1^-/-^Ifnlr1^-/-^*, *Cd40l^-/-^*, and *Tcra^-/-^* mice, models in which the thymic maturation of various APCs is altered as described above (**Fig. 5A, B, C**). Interestingly, the number of tDCs was not reduced in any of these models. This suggests that tDC do not require the same local signals (IFN and CD40L) that activate other thymic DC. Furthermore, thymic tDCs exhibited lover levels of IFN response and decreased Mx1^eGFP^ expression compared to other thymic APC, suggesting their limited IFN sensing within the thymic medullary region (**Fig. S4B and 5D**). This finding raises the possibility that tDCs may correspond to a previously described population of CX3CR1^+^ DCs capable of migrating into the thymus from peripheral tissues. Indeed, prior studies indicated that thymic CX3CR1^+^ DCs increase in number after weaning (approximately 21 days of life), yet their origin and characteristics were not addressed (Zegarra-Ruiz et al., 2021). Our transcriptional and lineage-tracing data showed that thymic CX3CR1^+^ DC can be comprised of moDC, moMacs, and tDCs (**Fig. 2C, F and 4C, G**). Notably, monocytes and macrophages are abundant early in life but decline with age (**Fig. 1D**), while tDCs are completely absent at birth, begin to appear around day 21 and increase over the first seven weeks of life (**Fig. 5E, F**).

**Figure 5.**
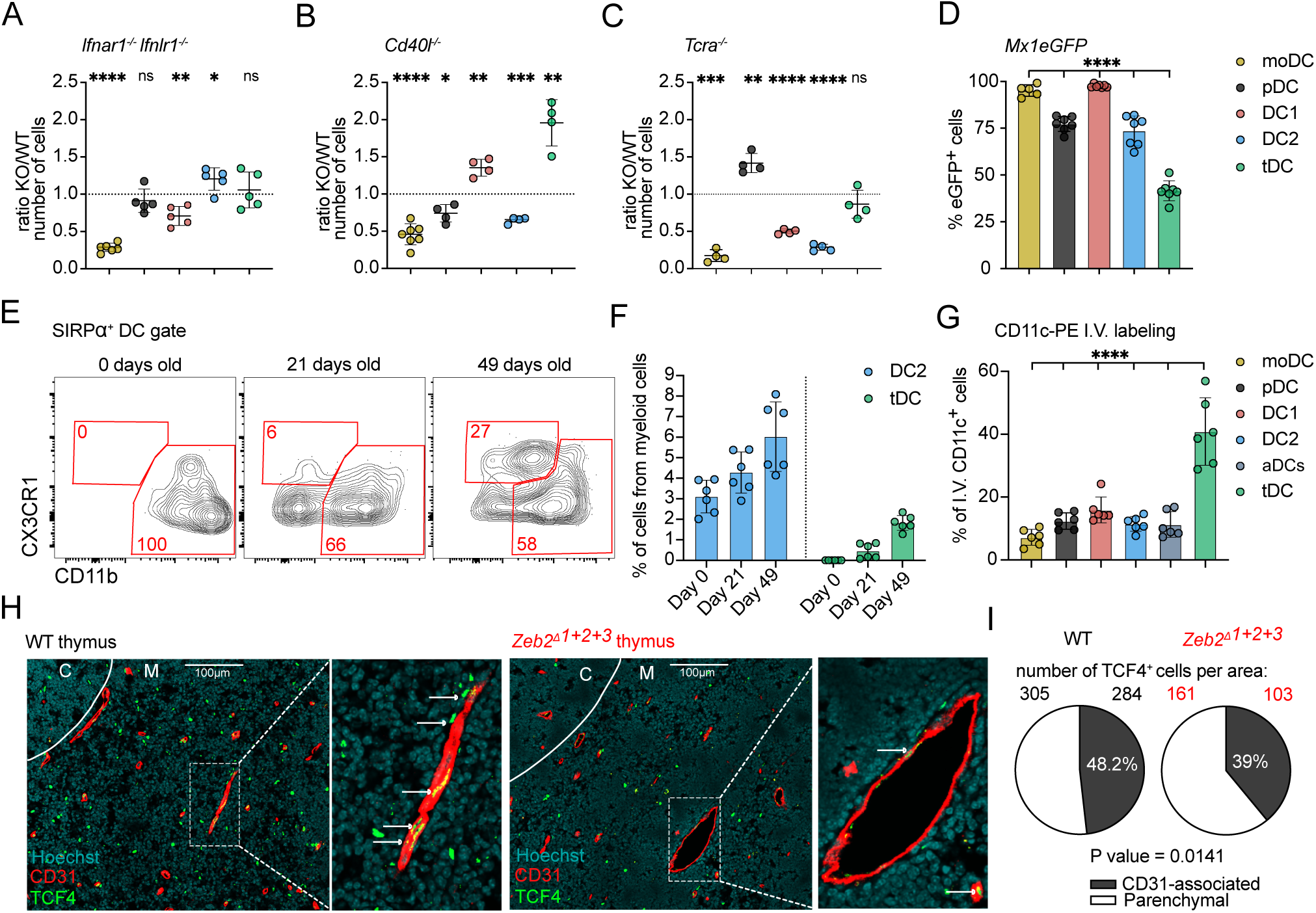
Transitional DC represent transendothelial cells. **(A)** Numbers of thymic monocyte-derived dendritic cells (moDC) and DCs (gated as in Fig. S4B) in *Ifnar1^-/-^* and *Ifnlr1^-/-^*mice, shown as KO/WT ratio of cell numbers, *n* = 5-6 mice from 2 independent experiments. **(B)** Numbers of thymic moDC and DCs in *Cd40l^-/-^*mice, shown as KO/WT ratio of cell numbers, *n* = 4-7 mice from 2 independent experiments. **(C)** Numbers of thymic moDC and DCs in *Tcra^-/-^*mice, shown as KO/WT ratio of cell numbers, *n* = 4 mice from 2 independent experiments. **(D)** Frequency of eGFP^+^ thymic moDC and DCs in *Mx1^eGFP^* mice, *n* = 5-7 mice from 3 independent experiments. **(E)** Representative flow cytometry plots showing thymic SIRPα^+^ DCs (gated as shown in Fig. S4B) from birth (0 days) to 49 days old mice. **(F)** Frequency of DC2 and tDC among total CD11c^+^ and CD11b^+^ cells in the thymus of C57BL/6 mice from birth (0 days old) to 49 days old mice, *n* = 6 mice. **(G)** Frequency of labeled thymic moDC and DCs by intra venous (I.V.) administration of anti-CD11c-PE antibody. Mice were euthanized 2 minutes after administration, *n* = 6 mice per 3 independent experiments. The cells were gated as shown in Fig. S5C. **(H)** Representative confocal microscopy images comparing the localization of TCF4^+^ cells in the thymus of WT C57BL/6 and *Zeb2^Δ1+2+3^*mice. The association with thymic microvessels were assessed by colocalization of TCF4^+^ cells with CD31^+^ positivity. Medullary region was identified by Hoechst staining. Scale bars represent 100 μm. **(I)** Numbers of free and CD31-associated TCF4^+^ cells in the specific thymus area of WT C57BL/6 and *Zeb2^Δ1+2+3^* mice. Data are shown as mean ± SD. Statistical analysis was performed by a one-sample *t* test and Wilcoxon test with a theoretical mean of 1 (A, B, and C), one-way ANOVA with multiple comparisons analysis (D and G), and two-sided Fisher’s exact test (I), \**p*≤0.05, \*\**p*≤0.01, \*\*\**p*≤0.001, \*\*\*\**p*≤0.0001, n.s. = not significant.

Previous work identified a population of transendothelial DC residing near microvessels in the thymic medulla as a circulating migratory DC population that brings blood-borne antigens into the thymus (Vollmann et al., 2021). These cells exhibited conventional DC2 characteristics and expressed CX3CR1 (Vollmann et al., 2021). To determine whether thymic tDC correspond to this transendothelial population, we intravenously (I.V.) injected mice with anti-CD11c-PE mAb, euthanized them two minutes post-injection, and analyzed the thymic DC pool via flow cytometry. The results showed that thymic tDCs bound the anti-CD11c mAb much more efficiently that other thymic DC subsets, with over 40% of these cells stained by I.V. labeling (**Fig. 5G and S4C**). This finding suggests that a substantial portion of tDCs are exposed to the bloodstream. Furthermore, all I.V. – labeled tDCs also displayed positivity when stained *ex-vivo* with anti-CD11C-PE-Cy7 (**Fig. S4C**), further confirming their transendotelial phenotype (Vollmann et al., 2021).

To test if thymic tDCs are positioned near microvessels, we performed confocal microscopy on frozen thymic sections stained with antibodies against TCF4 and the endothelial marker CD31 (**Fig, 5H left panel**). Notably, nearly 50% of TCF4^+^ cells (which includes pDC and tDC) were in close contact with CD31^+^ endothelial cells, idicating their presence in the perivascular space. The remaining half of the TCF4^+^ cells were located within the thymic parenchyma (**Fig. 5H, I**), aligning with our I.V. labeling data, where approximately 40% of tDCs were labeled (**Fig. 5G**). To distinguish tDC from pDC, we quantified TCF4^+^ cells in *Zeb2^Δ1+2+3^* mice, which lack tDCs while retaining pDCs (**Fig. S4D**). Interestingly, perivascular (CD31-associated) were strongly reduced in *Zeb2^Δ1+2+3^* mice (**Fig. 5H and 5I**). This suggests that the majority of CD31-associated TCF4^+^ cells correspond to tDCs. Together, these findings demonstrate the previous unappreciated finding that thymus immigrating DCs are tDC, of shared developmental origin with pDC.

## Discussion

Central tolerance is a crucial mechanism that prevents autoimmunity by eliminating self-reactive T cells during thymic selection. The magnitude of central tolerance is shaped by the diversity of self-p:MHC complexes that developing thymocytes encounter on thymic APCs (Klein and Petrozziello, 2024). The self-peptidome displayed by different thymic APC subsets varies significantly between cell types and is influenced by factors such as developmental origin, maturation and activation state, and key molecules that drive APC differentiation within the thymus or peripheral tissues (Spencer et al., 2015; Kim et al., 2025; Canesso et al., 2024). Notably, thymic hematopoietic APC exhibit a high degree of heterogeneity, the extent and functional implications of which remain incompletely understood. Here, we employed high-resolution scRNA-seq, complemented by various lineage-tracing and lineage-defining mouse models, to characterize the origin and lineage identity of thymic myeloid cells. In particular, we explored the internal heterogeneity within SIRPα^+^ DCs and found that what was previously described as ”cDC2” is composed of four developmentally distinct lineages. These include monocyte-derived moDC and moMacs, conventional DC2 with their activated counterparts (aDC2), and tDCs, which share a developmental origin with pDCs.

In this study, we identified four major populations of *Mafb*-expressing cells of monocytes/macrophage origin (Wu et al., 2016) (**Fig. 2A, B, C**). Phenotypically, these two lineages can be distinguished based on the expression of Ly6C, CD64 and MERTK. Monocytes express high levels of Ly6C, whereas macrophages are Ly6C-negative but express high levels of CD64 and MERTK. Notably, both populations contain thymus-specific subsets that upregulate CD11c, MHCII and co-stimulatory molecules (**Fig. 2D, E and S2C**). Moreover, the presence of these cells in the thymus is highly dependent on type I and type III IFN signaling (**Fig. 2J**). These characteristics led to the hypothesis that these cells represent activated subsets of monocytes and macrophages(Ashby et al., 2024). However, the fate-mapping experiments using *Ms4a3^Cre^*ROSA26^tdTomato^ mouse model clearly attributed both CD11c^+^MHCII^+^ monocytes (moDC) and CD11c^+^MHCII^+^ macrophages (moMacs) to the monocyte lineage (**Fig. 2F**). This finding suggests that thymic macrophage subsets represent distinct lineage identities rather than different activation states (Liu et al., 2019). This aligns with previous study describing thymic macrophage heterogeneity, which identified two major populations: TIM4^+^ embryonic-derived tissue-resident macrophages and hematopoietic stem cell-derived CX3CR1^+^ macrophages (Zhou et al., 2022). We believe that these correspond to macrophages and moMacs described in this study. Additionally, we hypothesize that these subsets also differ functionally, as MHCII^Low^ TIM4^+^ macrophages reside predominantly in the thymic cortex, whereas moMacs are localized in thymic medulla, where they upregulate MHCII and co-stimulatory molecules under the influence of IFNs, making them highly specialized thymic APCs (Zhou et al., 2022). The identification of moDC and moMacs that resemble conventional DC2 through the upregulation of CD11c, MHCII and co-stimulatory molecules provides valuable insight into the thymic APC subsets that are functionally equipped to induce T cell tolerance. The detailed characterization and lineage identity of these monocyte-derived thymic APC, presented in this study, will facilitate further research exploring the specific role of these subtypes in T cell clonal deletion and/or Treg selection.

The thymic DC pool contains homeostatically activated subsets characterized by the overexpression of CCR7, MHCII and co-stimulatory molecules (Park et al., 2020; Oh et al., 2018; Ardouin et al., 2016). DC activation is a process in which DCs transition from immature antigen-capturing cells to fully activated APCs capable of highly efficient antigen presentation (Guermonprez et al., 2002; Maier et al., 2020). Here, we show that thymic aDCs represent a continuum of cells spanning from early activated populations, which express both CCR7 and DC1- or DC2-lineage-defining molecules such as XCR1 or SIRPα, to fully matured late aDCs that substantially downregulate transcriptional and protein characteristics of their respective DC lineage. Notably, the DC1 population exhibited a more pronounced activation affect, as the CCR7^+^XCR1^-^ SIRPα^-^ cells predominantly represent aDC1 (**Fig. 3D, E and S3E, F**). Our study provides evidence that both thymic aDC populations are activated subtypes of DC1 and DC2, respectively, as they show clear dependence on either DC1-lineage defining *Batf3^-/-^* mouse model or the DC2 depleting *Zeb2^Δ1+2+3^* mouse (**Fig. 3F**). Interestingly, despite their distinct origins, aDC1 and aDC2 cells share highly similar transcriptional profiles, suggesting that they undergo a universal DC maturation program, ultimately leading to the formation of a functionally convergent APCs within the thymus (Park et al., 2020). Notably, the depletion of one DC subset is compensated by an increase in the other, preserving the total number of thymic DCs in both *Batf3^-/-^* and *Zeb2^Δ1+2+^* mice (**Fig. 3F**) (Hildner et al., 2008; Liu et al., 2022). We hypothesize that this mutual compensation between the thymic DC1 and DC2 lineages is driven by the availability of vacant niches that would normally be occupied by either subset. This dynamic adjustment presents a significant challenge in studying the distinct functions of DC1 and DC2 cells in central tolerance as the remaining subset may occupy a similar thymic microenvironment, facilitating their specific role in T cells selection.

As described previously, several thymic microenvironmental signals regulate APC maturation within the thymus (Ashby et al., 2024; Oh et al., 2018; Breed et al., 2022). Here, we show that the activation of moDC, moMacs and DC1 is highly regulated by type I and type III IFN signaling, whereas the maturation of DC2 relies on CD40-CD40L signaling. Notably, the absence of SP thymocytes in *Tcra^-/-^* mice leads to a marked reduction in all activated thymic APC subtypes, including moDC, moMacs, aDC1, and aDC2 (**Fig. 2J and 3G**). This finding aligns with previous study highliting the importance of CD4SP thymocytes and CD40 signaling in thymic DC2 maturation, with a lesser role in DC1 maturation (Oh et al., 2018). These results underscore the role of SP thymocytes in thymic APC maturation, either by providing CD40L or by promoting thymic IFN production. However, the direct effect of SP thymocytes on thymic APC maturation remains unclear, as the presence of SP T cells and the ablation of CD40L signaling also affect medullary thymic epithelial cells, which are key producer of type I and type III IFN, as well as other cytokines and chemokines (Akiyama et al., 2008; Ashby et al., 2024). Thus, it remains to be determined whether SP T cell directly signal to thymic DC to promote their activation or whether they modulate the overall thymic microenvironment, leading to secondary effects on APC maturation.

Historically, DC1 were described as thymus-resident, originating from intrathymic differentiation, whereas DC2 were thought to migrate from the periphery as fully differentiated cells (Porritt et al., 2003; Bonasio et al., 2006). However, previous studies using mouse parabiosis or photoconvertible mouse models suggested then only a minority of DC2 cells possess the ability to migrate into the thymus (Breed et al., 2022; Zegarra-Ruiz et al., 2021). It has been observed that pDC can enter the thymus in a CCR9-dependent fashion and present model antigens to developing T cells (Hadeiba et al., 2012). More recent research identified a population of transendothelial DC, localized near microvessels, enabling the transfer and presentation of blood-born antigens to developing T cells. The transendothelial positioning of these cells depends on CX3CR1 signaling (Vollmann et al., 2021). Additionally, a population of CX3CR1^+^ DCs has been observed to migrate into the thymus early in life, inducing the expansion of microbiota-specific T cells (Zegarra-Ruiz et al., 2021). Here we showed that the thymus accommodates a unique population of tDCs, phenotypically defined as TCF4^+^CX3CR1^+^CD14^+^SiglecH^-^CD11b^Low^ cells, which share a developmental origin with pDCs (**Fig. 4C-G**). These cells efficiently bound I.V. injected CD11c-PE antibody, were localized near thymic microvessels, and were present in the thymus only at later time points after the mouse weaning period (**Fig. 5E-G**). These characteristics clearly suggest that the descried tDCs represent thymus-immigrating and transendothelial DCs. Furthermore, based on their characteristics–such as their shared origin with pDCs and higher MHCII expression compared to thymic pDCs–we hypothesized that they may also represent the previously described thymus-immigrating “pDCs” responsible for model blood-born antigen presentation (Hadeiba et al., 2012). However, further investigation is required to confirm this hypothesis.

The exact role of DC2 populations in T cell selection remains unresolved, primarily due to the lack of comprehensive genetic tools that allow specific targeting of DC2. However, several studies utilizing partial DC2 depletion or model antigen presentation restricted to DC2 subsets suggest that thymic DC2 populations are more specialized for T cell clonal deletion rather than Treg selection (Breed et al., 2022; Bonasio et al., 2006; Vollmann et al., 2021). Previously we demonstrated that a substantial proportion of DC2 cells express CD301b lectin and that ablation of these cells significantly impairs the CD4^+^SP deletion (Breed et al., 2022). Notably, both DC2 and tDCs subpopulations express CD301b (**Fig. 4C**), making it difficult to distinguish functional differences between these populations. Here we provide evidence that the thymic DC2 lineage comprises two major, developmentally unrelated populations, allowing for the specific targeting of one subset to clarify its unique role in thymic T cell selection.

## Materials and methods

### Mice

Five- to eight-weeks old (unless otherwise states) male and female age-matched mice were used for experiments. Mice were housed in a specific pathogen-free facility under 12-h light: dark cycle at 22 ± 2 °C. C57BL/6J-*Ms4a3^em2(cre)Fgnx^*/J (*Ms4a3^Cre^*), B6.Cg-*Gt(ROSA)26Sor^tm14(CAG-tdTomato)Hze^*/J (*ROSA26^tdTomato^*), B6.129S2-*Ifnar1^tm1Agt^*/Mmjax (*Ifnar1^-/-^),* B6.129S2-*Cd40lg^tm1Imx^*/J (*Cd40l^-/-^*), B6.129S2-*Tcra^tm1Mom^*/J (*Tcra^-/-^*), B6(129S4)-*Xcr1^tm1.1(cre)Kmm^*/J (*Xcr1^Cre^*), B6.129X1-*Gt(ROSA)26Sor^tm1(EYFP)Cos^*/J (*ROSA26^eYFP^*), B6.129S(C)-*Batf3^tm1Kmm^*/J (*Batf3^-/-^*), and B6.Cg-Tg(Itgax-cre)1-1Reiz/J (*Itgax^Cre^*) were purchased from Jackson Laboratories. *Zeb2^Δ1+2+3^* mice (Liu et al., 2022) were kindly provided by K. Murphy (Washington University in St. Louise). *Ifnlr1^tm1.2Svko^* (*Ifnlr1^-/-^*) mice (Lin et al., 2016) were kindly provided by S. V. Kotenko (Rutgers New Jersey Medical School). B6.Cg-Mx1*^tm1.1Agsa^*/J (*Mx1^eGFP^*) mice (Uccellini and García-Sastre, 2018) were kindly provided by A. García-Sastre (Icahn School of Medicine at Mount Sinai). *Tcf4^fl/fl^* mice (Cisse et al., 2008) we kindly provided by B. Reizis (New York University). B6.Cg-Tg(CD2-icre)4Kio/J (*hCD2^iCre^*) were kindly provided by J. Idoyaga (University of California San Diego). All animal experiments were approved by the Institutional Animal Care and Use Committee of University of Minnesota.

### Cell isolation and flow cytometry

Thymic and splenic myeloid cells and B cells were isolated using Collagenase D (1mg/ml, Roche) dissolved in Hank’s balanced salt solution (HBSS) containing 2% fetal bovine serum (FBS), 10 mM HEPES, and Ca^2+^Mg^2+^ ions. Tissues were finely minced in 900μl of Collagenase D solution and incubated at 37°C for 15 minutes. The suspension was then pipetted up and down several times before undergoing a second incubation at 37°C for 20 minutes. After enzymatic digestion, the cell suspension was passed through 70 μm cell strainers, and the reaction was stopped by adding PBS with 2% FBS and 2 mM EDTA. Red blood cells from the thymus and spleen were lysed using ACK Lysis buffer (prepared in-house). For surface staining, cells were first incubated with an Fc block (anti-CD16/ CD32; 2.4G2, Tonbo Biosciences) for 15 min at 4°C. This was followed by a 30-minute incubation at 37°C with an anti-CCR7 antibody (4B12, Thermo Fisher Scientific). After washing, cells were further stained for 30 minutes at 4°C with the indicated surface antibodies. For intracellular TCF4 staining, cells were fixed with FoxP3 Transcription Factor Fix/Perm buffer (Thermo Fisher Scientific) for 2 hours and stained for 30 minutes in 1x Permeabilization buffer (Thermo Fisher Scientific) at 4°C. Samples were acquire with a BD Fortessa X-20, BD Fortessa H1770 or Cytec Aurora U1405 and analyzed with FlowJo v.10.10 (FlowJO LLC.).

### Antibodies

Antibodies purchased from BioLegend were the following: CD11c (N418), CD11b (M1/70), XCR1 (ZET), CX3CR1 (SA011F11), CD64 (X54-5/7.1), I-A/I-E (M5/114.15.2), CD14 (Sa14-2), CD172a (P84), CD88 (20/70), CD274 (10F.9G2), CD86 (A17199A), CD26 (H194-112), MERTK (2B10C42), VCAM-1 (429), TIM4 (RMT4-54). Antibodies purchased from BD Biosciences were the following: Siglec-F (E50-2440), Ly-6G (1AB), CD45R/B220 (RA3-6B2), Siglec-H (440c), CD197 (4B12), CD40 (3/23). Antibodies purchased from TONBO biosciences were the following: CD80 (16-10A1), F4/80 (BM8.1). Antibodies purchased from Thermo Fisher Scientific were the following: Ly-6C (HK1.40). Antibodies purchased from abcam were the following: TCF-4 (NCI-R159-6).

### Single-cell RNA sequencing and analysis

Sequencing and initial analysis were done at the University of Minnesota Genomics Center of the University of Minnesota as described previously (Breed et al., 2022). Thymic myeloid cells were isolated as described above and cells were MACS enriched for CD90.2^-^ cells, to deplete lymphocytes. CD11c/CD11b^+^ cells were FACS sorted and captured using the 10X Genomics 3′Single Cell v.3 chemistry platform and sequenced in a NovaSeq instrument. Prior to sequencing the quality control was assessed by Illumina-basicQC. Raw count data were loaded into R (v.4.4.1) and analyzed with the Seurat R package (v.5.1.0) (https://satijalab.org/seurat/). The dataset originally contained cells from multiple conditions identified by hashtag oligonucleotide (HTO) labeling. The Seurat function “HTODemux” was used to identify ‘doublet’ cells. After filtering out doublets, only C57BL/6J WT cells were selected for subsequent analysis. The mRNA expression data were then normalized using a log normalization method, where gene expression counts were normalized and scaled to correct for differences in sequencing depth and technical noise. To identify the most informative genes for downstream analysis, the “FindVariableFeatures” function in Seurat was used to select 2,000 highly variable genes. These features were then used for subsequent analyses, ensuring robust identification of cell clusters and states. Dimensionality reduction was performed using the “RunPCA” function. The top principal components were used as input for “RunUMAP”, which generated a two-dimensional visualization of the data based on the Uniform Manifold Approximation and Projection (UMAP) algorithm. Cell clustering was performed using the “FindClusters” function in Seurat, which applies a shared nearest neighbor (SNN)-based clustering approach. To visualize the clustering results the “DimPlot” function was used, which represents cells in UMAP space and colors them according to their assigned clusters. Gene expression patterns across clusters were visualized using “FeaturePlot”, which overlays expression levels of individual genes onto the UMAP projection. To identify differentially expressed genes (DEGs) between clusters, we applied the “FindMarkers” function, which performs statistical testing (Wilcoxon’s rank-sum test) to detect genes with significant expression differences between cell populations. These analyses and visualizations were conducted using R packages including, Seurat (v.5.1.0), ggplot2 (v.3.5.1), dplyr (v.1.1.4), and SeuratObject (v.5.0.2).

### Intravascular labeling

Intravascular labeling of thymic cells was done as described previously. Cells were labeled by intra venous injection (I.V.) of 1μg of PE-conjugated anti-CD11c mAb (clone N418). Mice were euthanized by CO_2_ asphyxiation followed by cervical dislocation 2 minutes post mAb injection. Isotype-PE antibody I.V. injection was used as control. Thymic myeloid cells were then isolated and analyzed as described above.

### Immunofluorescence

Thymi from C57BL/6J WT and were fixed in Cytofix/Cytoperm (BD Biosciences) at 4°C for 24 hours, followed by two washes in PBS. The tissues were then incubated in 30% sucrose at 4°C for 24 hours for cryoprotection. Afterward, the thymi were embedded in OCT compound (Sakura Finetek), frozen in the vapor phase of liquid nitrogen, and stored at −80°C until further processing. For analysis, frozen sections were dried overnight, rehydrated in PBS for 5 minutes, and blocked at room temperature for 60 minutes in PBS containing 0.3% Triton X-100, 1% BSA, and mouse Fc block. Sections were stained with primary antibodies TCF4 and CD31 (MEC13.3) overnight at 4°C. Sections were washed three times with PBS and stained with Hoechst 33342 (Thermo Fisher Scientific) for 5 min at room temperature. Sections were washed and mounted in ProLong Gold Antifade. Images were acquired using a Stellaris 8 confocal microscope (Leica Microsystems) and analyzed using Fiji and QuPath. The numbers of TCF4^+^ cells were calculated manually within the area of 1.7 mm^2^.

### Statistical analysis

For comparison of three or more datasets, ordinary one-way analysis of variance (ANOVA) with multiple comparisons test was used. One-sample t test and Wilcoxon test were used to perform single-column statistics. Two-sided Fisher’s exact test was used to analyze the differences in cell localization in microscopic images. Wilcoxon’s rank-sum test was used to identify DEGs in scRNA sequencing. P < 0.05 was considered significant. Sample size, experimental replicates, and additional details are provided in the figure legends. Statistical analyses were performed using GraphPad Prism 9.0.

## Data availability

scRNA-seq data are available in the NCBI’s GEO (https://www.ncbi.nlm.nih.gov/geo/) under accession number GSE198247. The main data supporting the findings of the present study are available in the article’s Supplementary Figures. Data are available from the corresponding authors upon request.

## Acknowledgements.

We thank A. García-Sastre (Icahn School of Medicine at Mount Sinai) for providing the *Mx1^eGFP^*, S. V. Kotenko (Rutgers New Jersey Medical School) for providing *Ifnlr1^-/-^*mice, K. Murphy (Washington University in St. Louise) for providing *Zeb2^Δ1+2+3^* mice and J. Ding and T. Dalzell for technical assistance, J. Motl (University Flow Cytometry Resource) for cell sorting, C. H. O’Connor and E. Stanley (University of Minnesota Genomics Center) for assistance with scRNA sequencing, and the University of Minnesota Research Animal Resources for animal husbandry. This work was supported by the National Institutes of Health Grant PO1 AI035296 and RO1 AI179749. Matouš Vobořil was a Cancer Research Institute Irvington fellow supported by the Cancer Research Institute (CRI4536).

## Authors contribution

M.V. and K.A.H. conceptualized and designed experiments and wrote the manuscript. M.V. performed majority of the experiments, with the help from K.M.A. and M.M.V. F.B.S. analyzed *hCD2^eYFP^*and *Itgax^Cre^Tcf4^fl/fl^* mice. R.J.M. generated microscopic experiments and help with scRNA seq analysis. J.I. provided *hCD2^eYFP^*and *Itgax^Cre^Tcf4^fl/fl^* mice and help to co-design some experiments. K.A.H. obtained funding for project. All authors provided feedback and approved the manuscript.

## Competing interests

J.I. reported personal fees from Immunitas Therapeutics outside the submitted work. The other authors declare no competing interests.

