## Supplemental material for "Thymic DC2 are heterogenous and include a novel population of transitional dendritic cells"

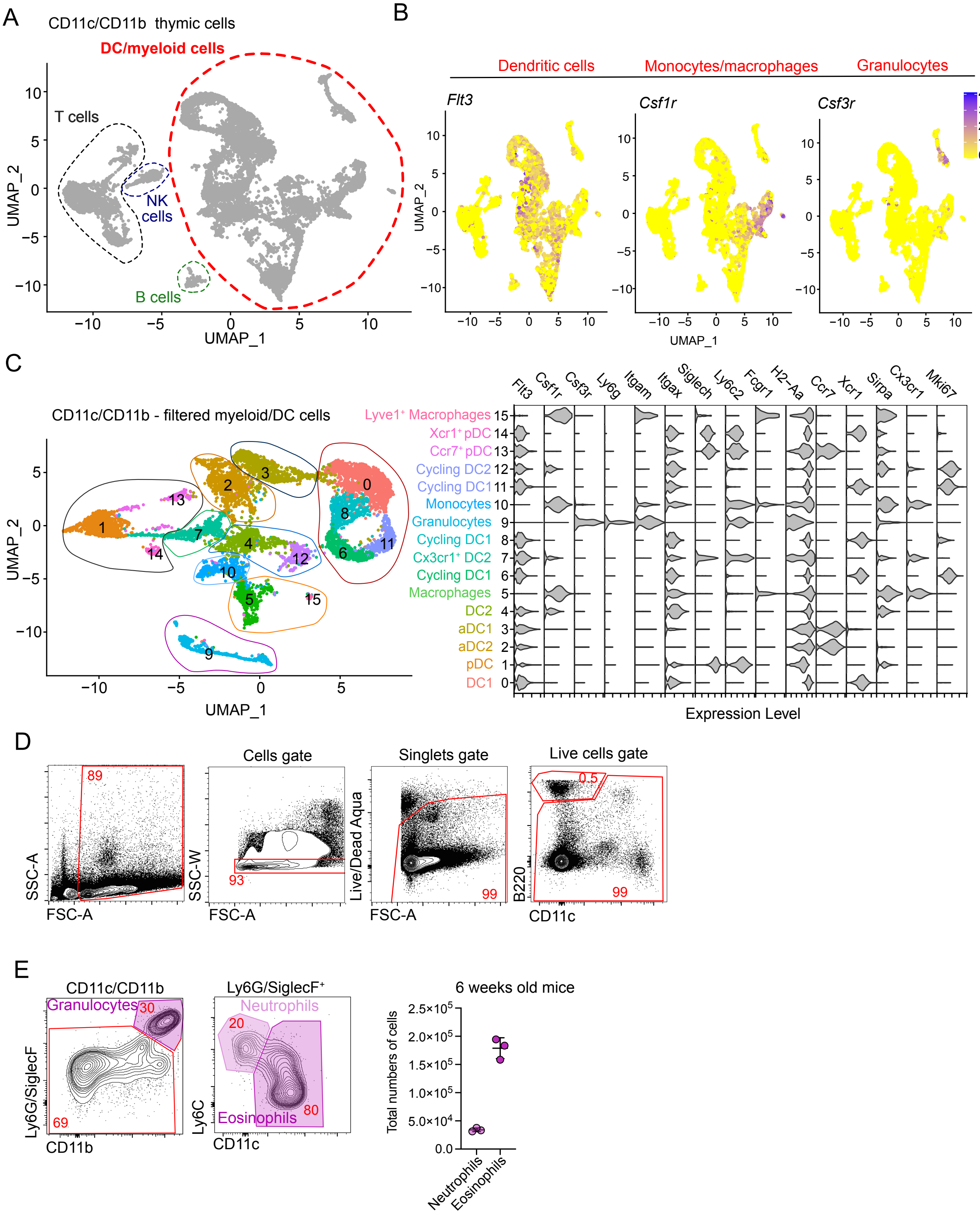

A
CD11c/CD11b thymic cells
DC/myeloid cells
T cells
NK cells
B cells
UMAP_1
UMAP_2
B
Dendritic cells
Monocytes/macrophages
Granulocytes
Flt3
Csf1r
Csf3r
UMAP_1
UMAP_2
C
CD11c/CD11b - filtered myeloid/DC cells
Flt3
Csf1r
Csf3r
Ly6g
Itgam
Itgax
Siglech
Ly6c2
Fcgr1
H2-Aa
Ccr7
Xcr1
Sirpa
Cx3cr1
Mki67
Lyve1+ Macrophages 15
Xcr1+ pDC 14
Ccr7+ pDC 13
Cycling DC2 12
Cycling DC1 11
Monocytes 10
Granulocytes 9
Cycling DC1 8
Cx3cr1+ DC2 7
Cycling DC1 6
Macrophages 5
DC2 4
aDC1 3
aDC2 2
pDC 1
DC1 0
Expression Level
UMAP_1
UMAP_2
D
Cells gate
Singlets gate
Live cells gate
89
93
99
0.5
99
SSC-A
FSC-A
SSC-W
FSC-A
Live/Dead Aqua
FSC-A
B220
CD11c
E
CD11c/CD11b
Granulocytes
30
69
Ly6G/SiglecF
CD11b
Ly6G/SiglecF+
Neutrophils
20
Eosinophils
80
Ly6C
CD11c
6 weeks old mice
Total numbers of cells
Neutrophils
Eosinophils

**Supplementary Figure 1. Single-cell RNA sequencing reveals heterogeneity in thymic dendritic cells.** **(A)** Single-cell RNA sequencing of CD11c$^{+}$ and CD11b$^{+}$ FACS-sorted cells from thymus of 7-week-old C57BL/6 mice. UMAP plots show the analysis of 11,586 transcriptome events, with dashed line representing clusters expressing *Flt3*, *Csf1r*, and *Csf3r*. **(B)** Feature plots showing the normalized expression of *Flt3*, *Csf1r*, and *Csf3r* in the clusters defined in (A). **(C)** UMAP plots show the analysis of 8,514 transcriptome events and identify 16 clusters of thymic myeloid cells. Violin plots show the normalized expression of signature genes in these clusters. **(D)** Representative flow cytometry gating strategy for pre-gating thymic myeloid cells. **(E)** Representative gating strategy for identifying thymic neutrophils (Ly6G/SiglecF$^{+}$Ly6C$^{+}$CD11c$^{-}$) and eosinophils (Ly6G/SiglecF$^{+}$Ly6C$^{-}$CD11c$^{+}$). The graph shows the total numbers of neutrophils and eosinophils per thymus in 7 weeks old C57BL/6 mice, $n$ = 3 mice. Data are shown as mean ± SD.

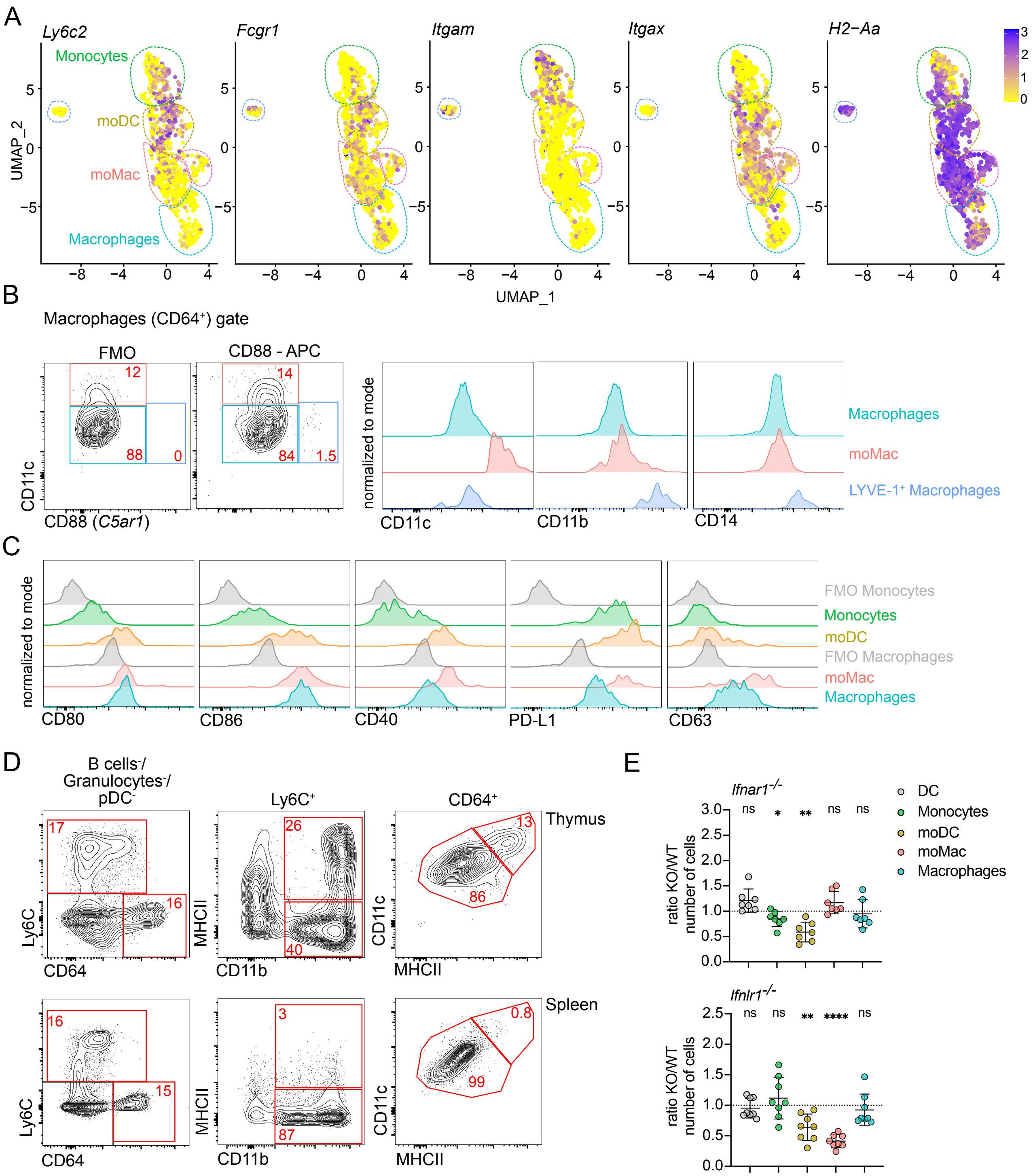
A
Ly6c2
Fcgr1
Itgam
Itgax
H2−Aa
Monocytes
moDC
moMac
Macrophages
UMAP_2
UMAP_1
B
Macrophages (CD64⁺) gate
FMO
CD88 - APC
12
88
0
14
84
1.5
CD11c
CD88 (C5ar1)
normalized to mode
CD11c
CD11b
CD14
Macrophages
moMac
LYVE-1⁺ Macrophages
C
normalized to mode
CD80
CD86
CD40
PD-L1
CD63
FMO Monocytes
Monocytes
moDC
FMO Macrophages
moMac
Macrophages
D
B cells⁻/
Granulocytes⁻/
pDC⁻
Ly6C⁺
CD64⁺
Thymus
17
16
26
40
13
86
Ly6C
CD64
MHCII
CD11b
CD11c
MHCII
Spleen
16
15
3
87
0.8
99
E
Ifnar1⁻/⁻
ns
*
**
ns
ns
ratio KO/WT
number of cells
DC
Monocytes
moDC
moMac
Macrophages
Ifnlr1⁻/⁻
ns
ns
**
****
ns

**Supplementary Figure 2. The thymus contains interferon-activated populations of monocyte-derived DC and macrophages. (A)** Feature plot showing the normalized expression of *Ly6c2*, *Fcgr1*, *Itgam*, *Itgax*, and *H2-Aa* in clusters identified in Fig. 2B. **(B)** Representative flow cytometry plots identifying LYVE-1$^{+}$ macrophages using CD88, CD11c, CD11b, and CD14 antibody staining. **(C)** Representative flow cytometry plots showing the normalized expression of CD80, CD86, CD40, PD-L1, and CD63, in thymic dendritic cells (DC; Ly6C$^{-}$CD64$^{-}$CD11c$^{+}$MHCII$^{+}$) and thymic monocyte and macrophage populations described in Fig 2D. **(D)** Representative flow cytometry plots comparing the monocyte and macrophages populations between thymus and spleen. **(E)** Numbers of thymic cells (gated as in D) in *Ifnar1*$^{-/-}$ and *Ifnlr1*$^{-/-}$ mice, shown as KO/WT ratio of cell numbers, $n$ = 7-8 mice from at least 2 independent experiments. Data are shown as mean ± SD. Statistical analysis was performed by a one-sample $t$ test and Wilcoxon test with a theoretical mean of 1, *$p \leq 0.05$, **$p \leq 0.01$, ****$p \leq 0.0001$, n.s. = not significant.

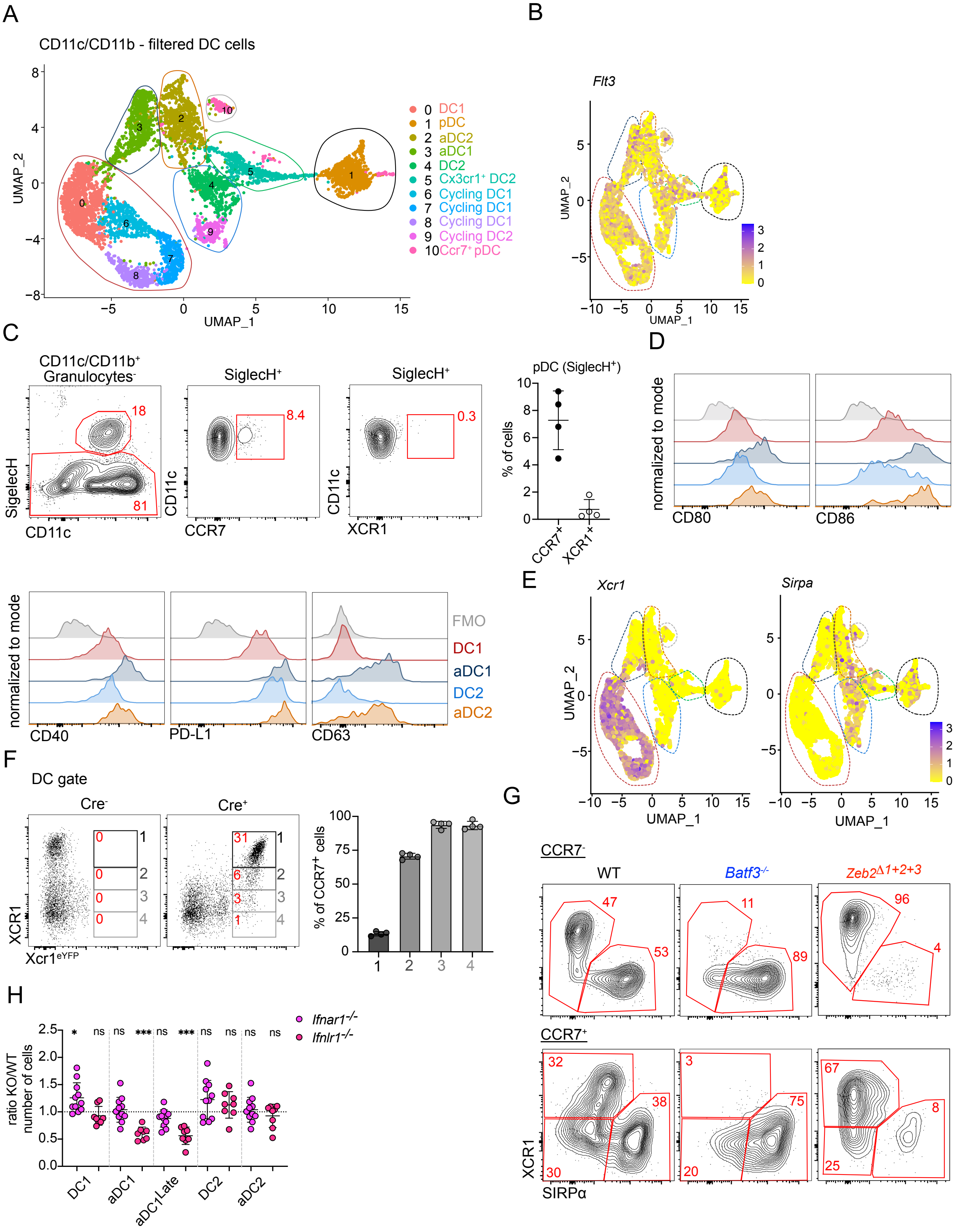
A
CD11c/CD11b - filtered DC cells
UMAP_1
UMAP_2
0 DC1
1 pDC
2 aDC2
3 aDC1
4 DC2
5 Cx3cr1+ DC2
6 Cycling DC1
7 Cycling DC1
8 Cycling DC1
9 Cycling DC2
10 Ccr7+ pDC
B
Flt3
C
CD11c/CD11b+ Granulocytes-
SiglecH+
SiglecH
CD11c
CCR7
XCR1
18
81
8.4
0.3
pDC (SiglecH+)
% of cells
CCR7+
XCR1+
D
normalized to mode
CD80
CD86
CD40
PD-L1
CD63
FMO
DC1
aDC1
DC2
aDC2
E
Xcr1
Sirpa
F
DC gate
Cre-
Cre+
XCR1
Xcr1eYFP
% of CCR7+ cells
G
CCR7-
CCR7+
WT
Batf3-/-
Zeb2Δ1+2+3
XCR1
SIRPα
47
53
11
89
96
4
32
38
30
3
75
20
67
8
25
H
Ifnar1-/-
Ifnlr1-/-
ratio KO/WT number of cells
DC1
aDC1
aDC1Late
DC2
aDC2
ns
*
***

**Supplementary Figure 3. Activation of conventional DC1 and DC2 requires distinct signals. (A)** Single-cell RNA sequencing of CD11c$^{+}$ and CD11b$^{+}$ FACS-sorted cells from thymus of 7-week-old C57BL/6 mice. Cells were bioinformatically filtered to include only clusters expressing *Flt3*. UMAP plots show the analysis of 6,928 transcriptome events, identifying 11 clusters. **(B)** Feature plot showing the normalized expression of *Flt3* in clusters identified in Fig. 3A. **(C)** Representative flow cytometry plots identifying CCR7$^{+}$ and XCR1$^{+}$ pDCs in the thymus. The graph shows frequency of CCR7$^{+}$ and XCR1$^{+}$ cells within the thymic SiglecH$^{+}$ population, $n = 4$ mice. **(D)** Representative flow cytometry plots showing the normalized expression of CD80, CD86, CD40, PD-L1, and CD63, in thymic dendritic cells populations defined in Fig. 3D. **(E)** Feature plots showing the normalized expression of *Xcr1* and *Sirpa* in clusters identified in Fig. 3A. **(F)** Representative flow cytometry plots showing expression of XCR1 in eYFP$^{+}$ cells from $Xcr1^{iCre}Rosa26^{eYFP}$ ($Xcr1^{eYFP}$) mice. The graph shows frequency of CCR7$^{+}$ cells in cell populations defined by flow cytometry, $n = 4$ mice, from 2 independent experiments. **(G)** Representative flow cytometry plots comparing the thymic DCs (gated as in Fig. 3C) in $Batf3^{-/-}$ and $Zeb2^{\Delta 1+2+3}$ mice. **(H)** Numbers of thymic DCs (gated as in Fig. 3C) in $Ifnar1^{-/-}$ and $Ifnlr1^{-/-}$ mice, shown as KO/WT ratio of cell numbers, $n = 8$-11 mice from at least 3 independent experiments. Data are shown as mean ± SD. Statistical analysis was performed by a one-sample *t* test and Wilcoxon test with a theoretical mean of 1, $^{*}p \leq 0.05$, $^{***}p \leq 0.001$, n.s. = not significant.

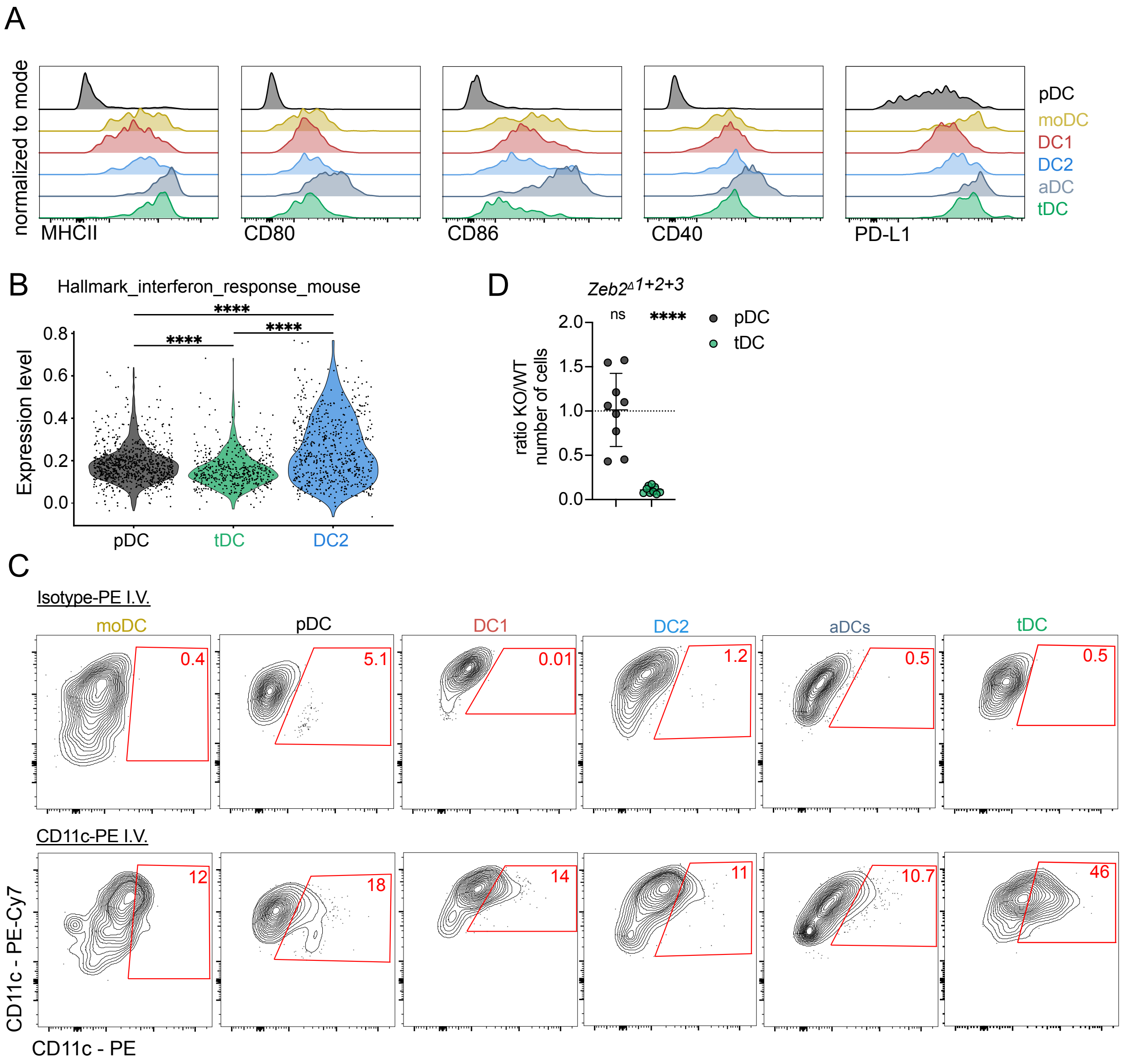


**Supplementary figure 4. Transitional DC represent transendothelial cells. (A)** Representative flow cytometry plots showing the normalized expression of MHCII, CD80, CD86, CD40, and PD-L1, in thymic dendritic cells populations gates as shown in Fig. 1C. **(B)** Violin plot displaying the normalized expression of gene associated with Hallmark interferon response mouse in clusters defined in Fig. 4B. **(C)** Numbers of thymic plasmacytoid DC (pDC) and transitional DC (tDC) in *Zeb2*$^{\Delta 1+2+3}$ mice, shown as KO/WT ratio of cell numbers, $n$ = 9 mice from 3 independent experiments. **(D)** Representative flow cytometry plots showing analysis of *ex-vivo* anti-CD11c-PE-Cy7 and intra venous (I.V.) anti-CD11c-PE labeling in thymic populations of monocytes-derived dendritic cells (moDC) and DCs. Data are shown as mean ± SD. Statistical analysis was performed by a Wilcoxon run-sum test (B) and one-sample $t$ test and Wilcoxon test with a theoretical mean of 1 (D), ****$p \leq 0.0001$, n.s. = not significant.
